# Function-associated scRNA-seq on single lung cancer organoids unravels the immune landscape of tumor parenchyma

**DOI:** 10.1101/2023.12.11.571047

**Authors:** Chang Liu, Kaiyi Li, Xizhao Sui, Tian Zhao, Ting Zhang, Zhongyao Chen, Hainan Wu, Chao Li, Hao Li, Fan Yang, Zhidong Liu, You-Yong Lu, Jun Wang, Xiaofang Chen, Peng Liu

## Abstract

In vitro models coupled with multimodal approaches are urgently needed to decipher the local tumor immune microenvironment (TIME) owing to the heterogeneous nature of immune cells and their diverse spatial distributions. Here we generate primary lung cancer organoids (pLCOs) by isolating the tumor cell clusters, including the infiltrated immune cells, from dissected lung cancer samples. A FascRNA-seq platform allowing both phenotypic evaluation and the scRNA-seq of all the single cells in an organoid was developed to dissect the TIME in individual pLCOs. Our analysis on 171 individual pLCOs derived from 7 patients revealed that pLCOs retained the fundamental features as well as intra-tumor heterogeneity of local TIME in the parenchyma of parental tumor tissues, providing a series of models with consistent genetic background but various TIME. Linking the single cell transcriptome data of individual pLCOs with their responses to ICB allowed us to confirm the pivotal role of CD8^+^ Ts in ICB induced anti-tumor immunity, to identify the potential tumor-reactive T cells with a set of 10 genes, and to unravel the factors regulating T cell activity.

## Introduction

Accurate in vitro models of the patient specific tumor immune microenvironment (TIME) and robust analysis techniques enabling in-depth dissection of the in vitro model are indispensable for the development of next generation immunotherapy. Imaging-based and sequencing-based analysis of the surgically dissected patient tissues has enabled the deconvolution of not only the cell types that constitute the TIME but also their cell-state heterogeneity (*1–4*). However, they can only provide the snapshots rather than the dynamics of TIME, i.e., the subtle changes of TIME in response to treatments. Simplified *in vitro* models representing the *in vivo* tumor micro-niche, retaining the local TIME, and facilitating the modulation of drug treatment are desired. Patient-derived organoids (PDOs) have been widely accepted as a valuable *in vitro* tumor model that faithfully recapitulates the histological and molecular characteristics of original tumors (*5–9*). Depending on the methods for organoid generation and the period of *in vitro* culture, tumor organoids can preserve the immune components of the parental tissues (*10, 11*). For example, the PDOs established using mechanical crushing methods and cultured either in a gel matrix (*12*) or an air-liquid interface (*13*) contained tumor infiltrating lymphocytes and responded to ICB (*12, 13*). However, whether they recapitulate the inter- and intra-patient heterogeneity of local TIME are to be proved. More importantly, limited by the number of cells, current analysis methods for PDOs are mainly phenotypic- or imaging-based, impeding the fully dissection and wide application of PDOs as a model of patient TIME.

Single cell RNA sequencing (scRNA-seq) is an indispensable tool to dissect the cellular diversity within a complex biological system. However, current sequencing-based techniques are not applicable for deciphering the TIME in an organoid owing to the disruption of the spatial distribution of immune cells. For example, the commonly used scRNA-seq techniques, such as 10x Chromium, stochastically capture and encode single cells, leading to the requirement of minimum 10^3^-10^4^ starting cells with a considerable cell loss (*14*). The efficient processing of every single cell from each individual organoid remains challenging. Although ample single cells can be digested from PDOs and pooled for processing, the link of single cells with their specific microenvironment within individual PDO will be disrupted (i.e., immune cells in individual organoids cannot be identified). Actually, the retrospection of single cells to their parental organoids is extremely important, as it would enable the single-cell data analysis under the supervision of phenotypic responses of organoids.

In order to probe the immune microenvironment in local tumor micro-niches, we established primary lung cancer organoids (pLCOs) from dissected tumor tissues of lung cancer patients. Owing to the nature of the mechanical processing methods, clusters of tumor cells were isolated and the infiltrating immune cells were retained. We proved that the pLCOs represent the general and patient specific features of local TIME in tumor parenchyma. To perform single cell analysis on individual pLCOs, we developed a function-associated scRNA-seq (FascRNA-seq) system, with which the phenotypic changes of pLCOs upon ICB treatment were evaluated, and at the meanwhile, the transcriptomes of all the cells from each pLCO were sequenced and correlated to the phenotypic data (Fig. 1A). Subsequent analysis certified that the anti-PD-1 (aPD-1)-induced cytotoxic effect was mediated by CD8^+^ T cells which showed patient specific activation patterns. Our methods can be used for characterizing the local tumor micro-niche, identifying the tumor-reactive T cells, and potentially facilitating the development of new immunotherapy strategies.

**Fig. 1.**
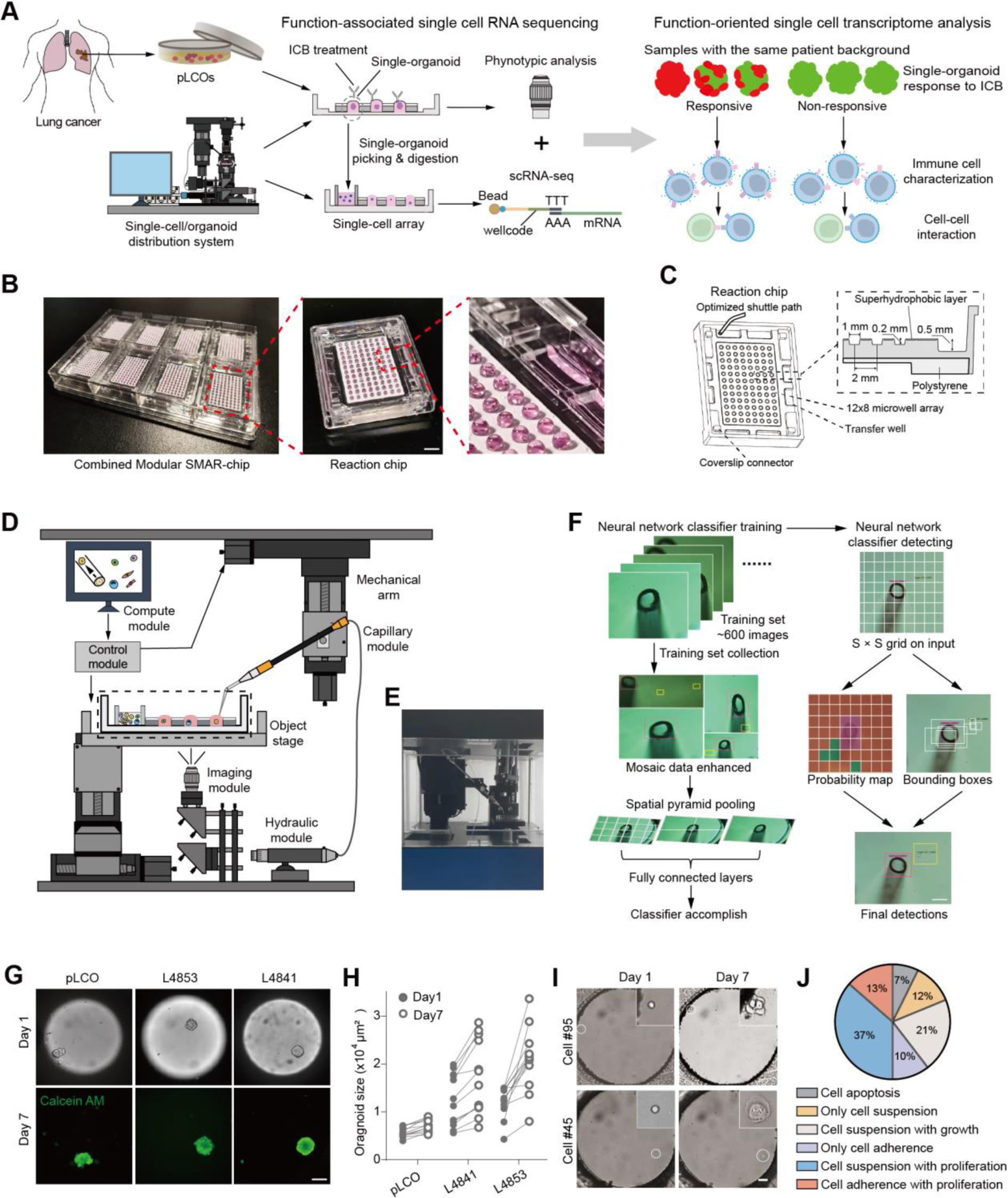
The single cell and single organoid distribution platform. **(A)** Schematic design of the single pLCO analysis using FascRNA-seq. After patient-derived lung cancer organoids were generated, single pLCOs were distributed into MoSMAR-chip microwells followed by 5 to 7-day aPD-1 or IgG treatment. Individual pLCO was stained with Calcein AM/PI for phenotypic evaluation, then digested into single cells which were subsequently distributed into another MoSMAR-chip for scRNA-seq. **(B)** Images of the MoSMAR-chip. 8 reaction chips were installed on the chip frame and each reaction chip consists of 12×8 microwells. Scale bar, 5 mm. **(C)** Schematic diagram of the reaction chip, showing the microwell array, sample transfer wells, and the connectors for placing transfer coverslip. **(D)** Schematic diagram and **(E)** image of the automated single-cell/organoid distribution system, consisting of the imaging, computing, motion control, and capillary modules. **(F)** Schematic diagram of the process for establishing the YOLO neural network for single cell/organoid recognition. The YOLO was trained with ∼600 collected images to form the classifier, then the neural network classifier was applied to identify capillary opening, single cells, and single organoids. **(G)** Images of single organoids distributed in the microwells and traced for 7 days. At day 7, the organoids were stained with Calcein/AM to confirm their viability. Scale bar: 100 μm. **(H)** Size of the traced single organoids at day1 and day 7 post distribution. **(I)** Images of single cells disgested from L4853 organoids and cultured in the microwells of a MoSMAR-chip. The circle indicates the single cell which is shown by the enlarged image at the upper right cornor. Scale bar, 50 μm. **(J)** Various circumstances of L4853 derived single cell culture after 7 days in the microwells of a MoSMAR-chip. (**** P<0.0001, paired student’s *t* test, n=96)

## Results

### Automated single-cell and single-organoid distributions

The cornerstones of the FascRNA-seq system are the modular superhydrophobic microwell array chip (MoSMAR-chip), which is the reactor for tumor organoid culture, drug sensitivity test, and scRNA-seq sample preparation (Fig. 1, B and C, fig. S1), and the automated single cell/organoid distribution instrument (SCDI), which automatically delivers single cells/organoids to each microwell on the MoSMAR-chip (Fig. 1, D to F, fig. S2). The MoSMAR-chip fabricated by injection molding with polystyrene consists of eight 96-microwell reaction chips assembled in a chip frame, achieving a higher throughput of 768 for FascRNA-seq. The reaction chip is designed to accommodate a transfer coverslip (Fig 1C), which can deliver reagents to each microwell by a “spot-cover” method. The droplet-to-droplet contact between the reaction chip and the transfer coverslip is realized due to the home-made superhydrophobic paint on the top surfaces of these chips (*15*).In the reaction chip, the 12×8 microwell array is surrounded by eight large transfer wells (Fig 1C), in which a large number of single cells can be preloaded or organoids can be digested to single cells for the subsequent single-cell distribution with minimum transfer distances.

We developed an automated single cell distribution instrument employing a microneedle-based liquid handling robotic under the control of an AI-assisted image processing and recognition software. The image recognition employing the YOLO neural network (*16*) computing framework was trained with a dataset of ∼600 images, achieving the real-time recognition and positioning of capillary needles and single cells (Fig 1F, fig. S2A). The cell distribution can be operated in different modes for rare (i.e., cells isolated from an organoid) or sufficient cell samples (i.e., digested cell line samples) (fig. S2B) and adapted to capillaries with different opening diameters for samples with different sizes (i.e., single cells and organoids). The image recognition accuracy 86.5% and 72.3%, and the distribution accuracy 91.3% and 94.6% were achieved for rare single-cell and single-organoid samples, respectively (fig. S2, C to F) (Movie S1 and 2). The compact layout of the MoSMAR-chip, the short distance between the transfer wells and the microwell array, and the barrier-free interface between microwells make the on-chip operation much more efficient than that on conventional multi-well plates (Movie S3). The single-cell distribution rate was significantly increased from 7.5 s/cell on plates to 2.5 s/cell on chips (fig. S2F). We delivered 2 organoid lines generated in our previous study and 1 primary organoid at day 1 post sample collection on the MoSMAR-chip. After 1-week culture, all the organoids had increased areas at day 7 compared to day1 (Fig. 1, G and H). In addition, single cells digested from organoid line L4853 (*15*) also showed good viability (Fig. 1, I and J), indicating the good biocompatibility of the distributing prosess.

### Function-associated scRNA-seq

To perform scRNA-seq on single cells digested form organoids, we upgraded our previously developed Group-seq methodology (*17*) to the single-cell level with a higher throughput. Since we need to record the barcode in each microwell to trace single cells, we employed a ligation-based barcode synthesis method to lower the costs of the long capture oligos that attach to magnetic beads for mRNA capture (Fig. 2A). To validate the performance of the on-chip scRNA-seq, 192 human lung cancer cells (H2122) were distributed and sequenced. On average, more than 30,000 transcripts corresponding to 5820 genes were detected from each cell, while the ratio of mitochondria genes is less than 11% (Fig. 2B), indicating the good library quality. The sequencing data had a mapping rate of ∼78%, an exon mapping rate of ∼63%, and a unique mapping rate of ∼96% (Fig. 2C), comparable to previously reported magnetic bead-based scRNA-seq methods, such as seq-well (*18*) and drop-seq (*19*). Furthermore, the average of the 192 scRNA-seq data had good correlation with the normal bulk seq results (Pearson correlation coefficient (PCC) r=0.93, Fig. 2D). In order to validate the repeatability, the experiments were repeated for 3 times and PCC of r>0.95 was obtained for any two repetitions (Fig. 2E). We also tested the inter-microwell cross contamination by delivering human cells (H2122) and mouse cells (NIH3T3) into alternate lines on the chip and found more than 99% of the 192 microwells were free of genes from the other species (Fig. 2F). At last, scRNA-seq of the mixture of H2122 and Huh7, a liver cancer cell line, revealed two separate groups featured with expression of marker genes of lung and liver cells, respectively (Fig. 2G). These data demonstrated the reliability of our MoSMAR-chip-based scRNA-seq.

**Fig. 2.**
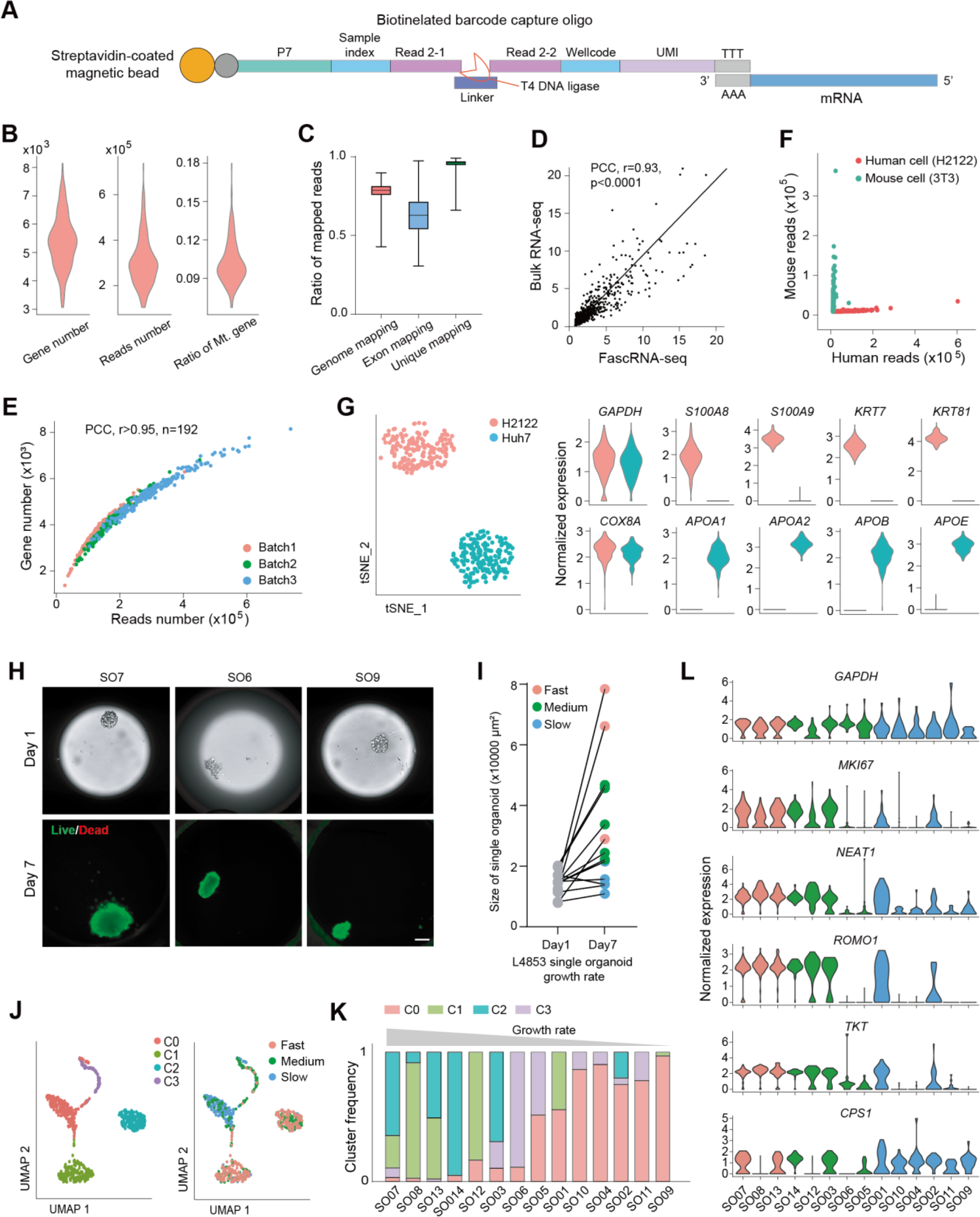
Verification of FascRNA-seq. **(A)** Structure of the 3’ mRNA capture oligo for FascRNA-seq(*17*). **(B)** Violin plots of the number of genes (average 5380), mapped reads (average 30522), and ratio of mitochondria-related genes (Mt. gene) detected by the on-chip scRNA-seq for 192 H2122 cells. **(C)** Ratio of mapped reads for H2122. The center line represents the median value. The bounds of box represent the median values of the upper half and the lower half. The bounds of whiskers represent the maxima and the minima. **(D)** Correlation of the average of the 192 single-cell transcriptomes with bulk RNA-seq data. The top 1000 genes detected in the on-chip scRNA-seq were compared. **(E)** Numbers of reads and genes detected in single cells for three repeats of the on-chip scRNA-seq. **(F)** Numbers of mouse reads and human reads detected in each microwell of a MoSMAR-chip which is distributed with single mouse cells (NIH3T3) and human cells (H2122) in alternate lines. **(G)** Cell type profiling of the H2122 and Huh7 cell lines using the on-chip scRNA-seq. Left: UMAP of the single-cell transcriptome data of the two types of cells. Right: Violin plots showing the expression levels of signature genes for lung (H2122) and liver (Huh7) cells. **(H)** Representative images of the 14 organoids of L4853 cultured on the chip for 7 days. On day 7, the organoids were stained with Calein AM/PI to verify their viability. Scale bar, 100 μm. **(I)** Quantification of the growth rates of the 14 organoids. Individual organoids were categorized into three groups according to their growth rates (GR) (Fast, GR ≥ 3; Medium, 1.5 < GR < 3; Slow, GR ≤ 1.5). **(J)** UMAP projections of 672 single cells derived from 14 organoids of L4853. Cells are color labeled with the unsupervised UMAP clusters (left) or the high, medium, and low categories according to the growth rates of the parental organoids. **(K)** Proportions of cells of the 4 UMAP clusters in the 14 organoids. **(L)** Average expression of selected reference genes (*GAPDH*), typical proliferation (*MKI67*) and malignancy related genes in single organoids. Note the upregulated expression of these genes in fast growing organoids. Individual organoids are color labeled with the high, medium, and low categories according to their growth rates.

Next, we went through the whole FascRNA-seq process (fig. S3A) using a previously established lung cancer organoid line, L4853 (*15*). A total of 14 organoids were seeded on the chip and traced for a week (Fig. 2H). The growth ratios (area of an organoid at day 7 divided by area at day 1) of these observed organoids varied from 1.04 to 5.29 (Fig 2I). Then, these 14 organoids were picked, digested into single cells (672 cells in total), distributed on the MoSMAR-chip and processed for scRNA-seq. Unlike the 10x Genomics platform where a single cell cannot be traced back to the parental organoids, the FascRNA-seq system can identify all the single cells from each individual organoid. Seurat analysis of the transcriptome data of the 672 cells from these 14 organoids resulted in 4 major clusters. C0 and C3 mainly consist of cells from slowly growing organoids while the majority of cells in C1 and C2 were from organoids with high growth rates (Fig. 2, J and K). Consistent with the phenotypic data, C1 and C2 had higher expression of genes related to proliferation and malignancy, including *ROMO1, MKI67, NEAT1* and *TKT* (Fig. 2L, fig. S3B). Pseudo time analysis of the potential developmental trajectories indicated that these genes were upregulated along the inferred developmental trajectory (fig. S3C), suggesting the trend towards increased malignancy during the long-term culture.

### pLCOs retain the local immune microenvironment in tumor parenchyma

In order to retain the TIME, we employed a digestion-free sample processing method developed previously (*15*), i.e., collecting the 40-100 µm cell aggregates produced by mechanically grinding the tumor samples. Unlike the enzymatic digestion, the cell-cell junctions were not interrupted during the processing (Fig. 3A). We have proved the enrichment of cancer cells and the exclusion of stromal cells, since the large ECM (extracellular matrix) fibers abundant in stroma cannot be broken by gentle grinding. Consistently, more organoids were generated from late-stage samples which usually had higher contents of neoplastic cell (*15*). Therefore, we speculate that the primary lung cancer organoids (pLCOs) generated by this method retain the local micro-niche in tumor parenchyma, including the TIME.

**Fig. 3.**
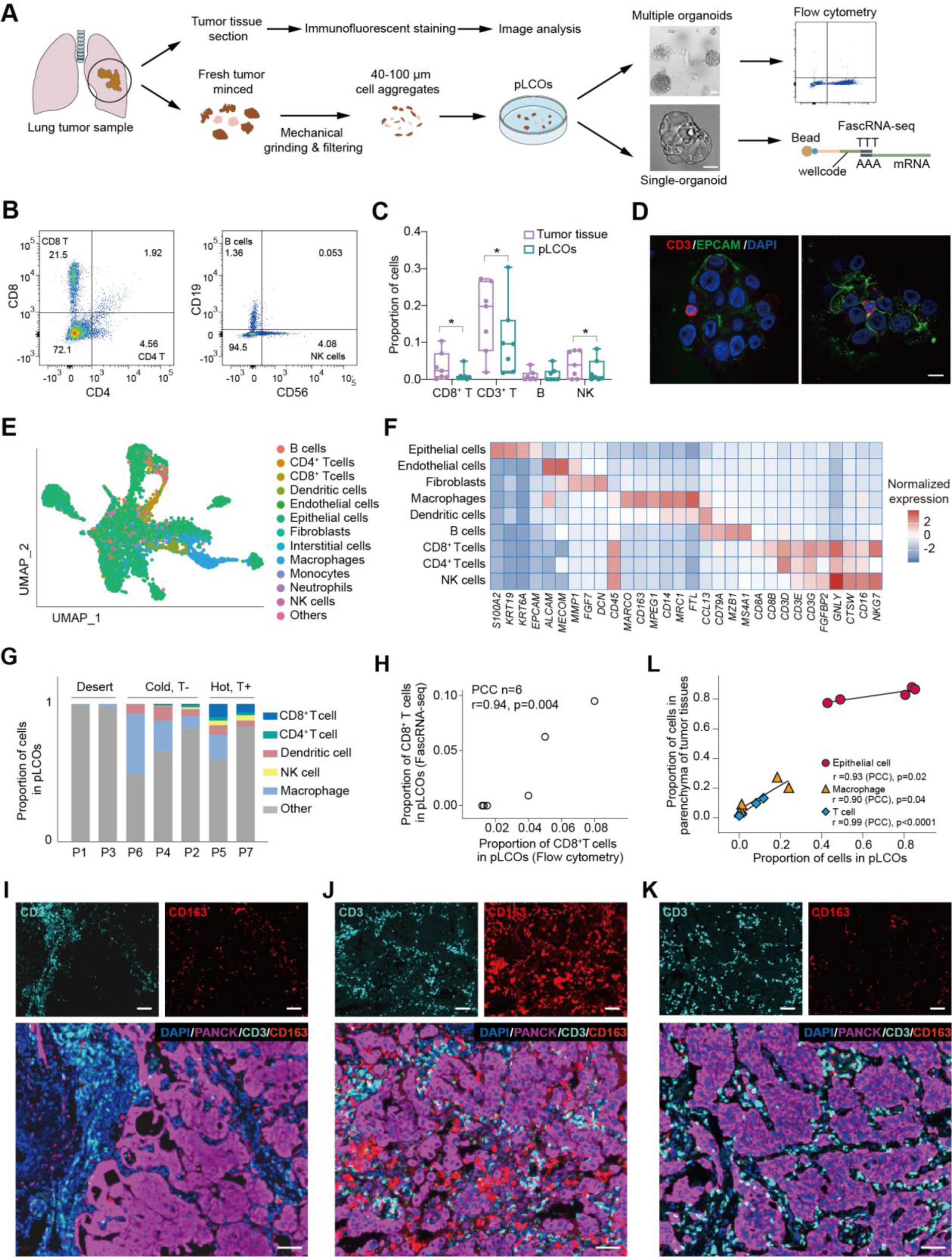
Characterization of primary lung cancer organoids (pLCOs) **(A)** Scheme for pLCO generation and characterization. The pLCOs were generated by mechanical grinding and filtering of lung cancer samples, characterized by flow cytometry, and FascRNA-seq. Scale bars, 50 μm. **(B)** Flow cytometry results showing the presence lymphocytes, including CD4^+^ T, CD8^+^ T, NK cells, and B cells in pLCOs. **(C)** Proportions of lymphocytes in pLCOs and the parental tumor tissues detected by flow cytometry. Note the significant decrease of T cells in pLCOs compared to tumor tissues. The center line represents the median value. The bounds of box represent the median values of the upper half and the lower half. The bounds of whiskers represent the maxima and the minima. (*P<0.05, paired student’s *t* tests, n=7). **(D)** Immunofluorescent staining images of the cryosection of pLCOs. Note the presence of CD3^+^ T cells which is in direct contact with the EpCAM+ epithelial cells. Scale bar, 10 μm. **(E)** UMAP plot of the 6240 single cells from 171 pLCOs. The single cells were color labeled with the annotated cell types. **(F)** Average expression levels of marker genes for the annotated cell types. **(G)** Proportions of the indicated immune cells in pLCOs derived from 7 lung cancer samples. Samples are categorized into three types according to the frequencies of the immune cells (desert, no immune cells; cold, few T cells; hot, abundant T cells). **(H)** Pearson correlation coefficient (PCC) analysis between the flow cytometry and the FascRNA-seq results on the proportion of CD8^+^ T cells in pLCOs derived from different patient samples. **(I to K)** Images of the immunofluorescent staining on the sections of tumor tissues P3 **(I)**, P5 **(J)** and P7 **(K)**. The tissue sections were stained with CD3, CD163, and PanCK to detect the T cells, macrophages, and cancer cells, respectively. Scale bars, 100 μm. **(L)** PCC analysis revealed significant correlation of cell proportions between pLCOs and the parenchyma region of tissue sections.

To prove this speculation, we first digested the organoids to single cells for flow cytometry analysis. Lymphocytes, including T (CD4^+^, CD8^+^), B (CD19^+^), and Natural killer (NK, CD56^+^) cells, were detected (Fig. 3B), although the proportions of T and NK in pLCOs were significantly lower than those in parental tumor tissues (Fig. 3C), probably due to the existence blood vessels and tertiary lymph nodes in the stroma. Immunofluorescent staining on frozen section further demonstrated the presence of T cells in the pLCOs, in contact with the surrounding EpCAM expressing cancer cells (Fig. 3D). Previous reports as well as our other studies indicated that the number of immune cells in tumor organoids decreased gradually with time in culture (*13*), so all our experiments were performed on pLCOs within 1 week since sample processing.

Next, FascRNA-seq was performed on individual pLCOs to probe the response to ICB and the single-cell transcription profiles. A total of 171 organoids generated from 7 patient samples were distributed on the MoSMAR-chip, treated with either anti-PD1 antibody (aPD-1) or control IgG for 7 days, and finally digested to single cells for scRNA-seq (Fig. 4A). The corresponding scRNA-seq data derived from individual pLCOs was displayed as a landscape under UMAP view (Fig. 3E, fig. S4A). The expression levels of marker genes were in good agreement with the annotated cell types (Fig. 3F). In addition, the proportion of CD8^+^ T cells derived from FascRNA-seq and flow cytometry analysis showed a good correlation, confirming the accuracy of the annotation (Fig. 3H). Consistent with the epithelial nature of the pLCOs (*20*), the majority part (64.3%) of the cells was epithelial and the copy number variations (CNVs) analysis based on the average expression profiles across chromosomal intervals revealed that most of the cells were malignant with genetic mutations (fig. S4, B and C). Importantly, 25.7% of sequenced cells were immune cells, including both myeloid and lymphoid cells. Marchophages (Mph), dendritic cells (DC), T cells, and NK cells were relatively abundant which made up 13.6%, 5.2%, 4.2%, and 1.4% of total cells, while B cells and monocytes were rare (0.58% and 0.66%), echoing their less importance in tumor immunity. pLCOs derived from the 7 patient samples represent the three major classes of TIME (*21*). P1 and P3 pLCOs show the feature of immune “desert” with rare immune cells. P2, P4, and P6 pLCOs represent T cell excluded “cold tumor”. P5 and P7 are “hot” with considerable amounts of T cells (Fig. 3G).

**Fig. 4.**
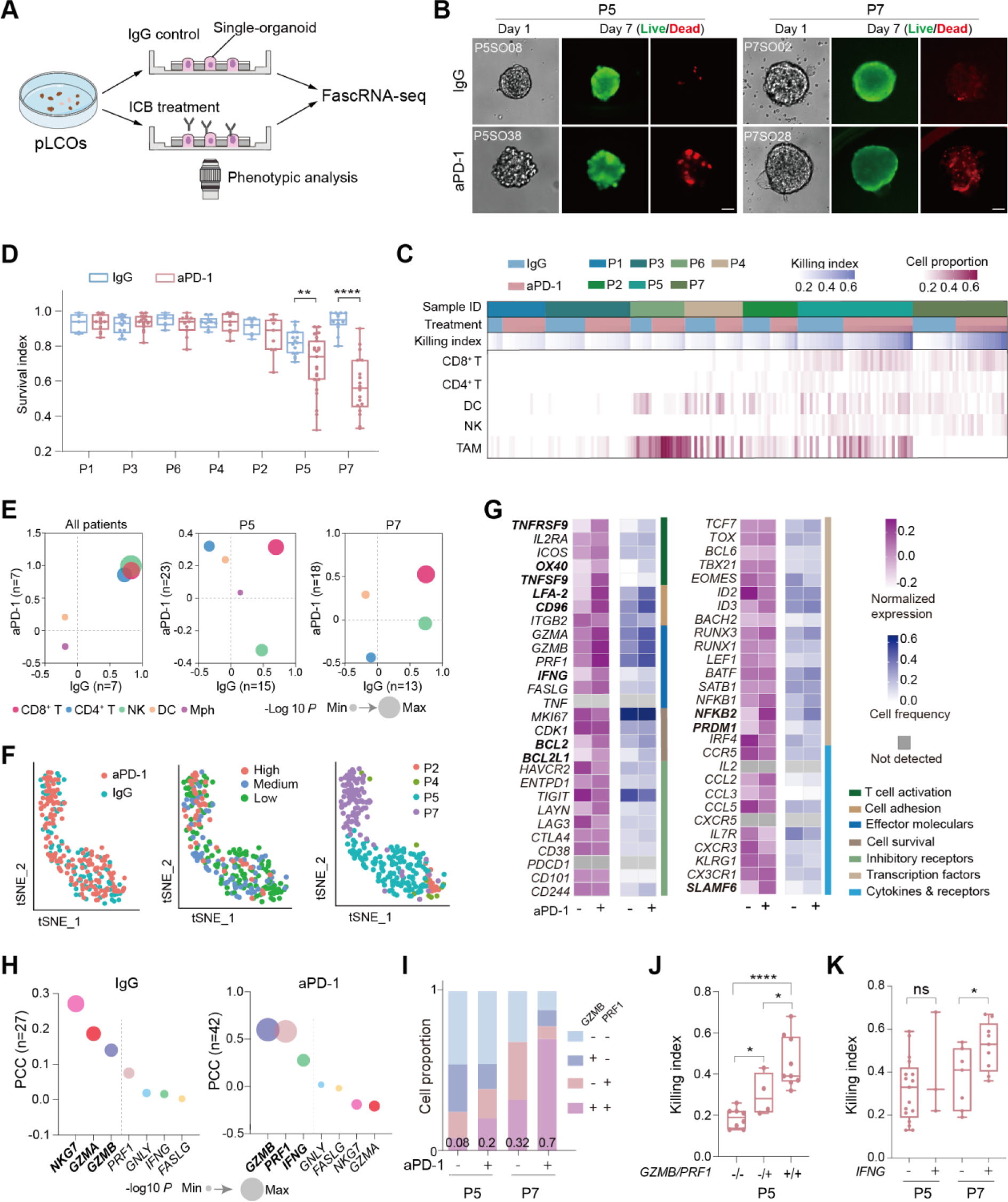
CD8^+^ Ts mediate the response of pLCOs to anti-PD-1 treatment. **(A)** Scheme of the drug treatment experiment. **(B)** Images of the individual pLCOs under IgG or aPD-1 treatments. pLCOs were stained with Calcein AM/PI on day 7. Scale bars, 50 μm. **(C)** Heatmap showing the proportions of immune cells and the quantification of killing index in the 171 single pLCOs with or without aPD-1 treatment. **(D)** Comparison of the survival of individual organoids under aPD-1 or control treatments. **(E)** PCC between immune cell proportion and the Killing index. The x and y axis show PCC for the aPD-1 and the control treatment conditions, respectively. The data points are color labeled by the types of immune cells. The diameter of the point represents the geometric mean of the P value. Left: PCC analysis at the patient level where the average of organoids from the same patient sample were used as the patient data. Middle and right: PCC at the pLCO level of P5 (middle) and P7 (right). **(F)** The t-SNE projection of the 191 CD8^+^ T cells from all the pLCOs. Data point of each cell was labeled by treatment conditions (left), patient (right) and the categories of killing index (Ki) (middle, High: Ki ≥ 0.5; Medium: 0.25 < Ki < 0.5; Low: Ki ≤ 0.25). **(G)** Heatmap showing the average of the normalized expression of T cell function related genes under the two treatment conditions and the proportion of cells with positive expression. The boldfaces indicate genes with significantly different levels under the two treatment conditions. **(H)** PCC between the average expression level of the effector molecules in single pLCOs and the killing index. **(I)** Quantification of the frequency of CD8^+^ T cells expressing 0, 1, or 2 of the effector molecules *GZMB* and *PRF1*. **(J)** Comparison of cell death in P5 pLCOs with CD8^+^ T cells expression 0, 1, or 2 of the molecules, *GZMB* and *PRF1*. **(K)** Comparison of cell death in P5 and P7 pLCOs with or without *IFNG* expressing CD8^+^ T cells. The center lines in the box plots of **(D, J and K)** represents the median values. The bounds of box represent the median values of the upper half and the lower half. The bounds of whiskers represent the maxima and the minima. (*P < 0.05, **P < 0.01, ***P < 0.001, ****P<0.0001, unpaired, two-sided student’s *t* tests).

To demonstrate whether the TIME in pLCOs represent the *in vivo* counterparts, we performed immunofluorescent staining on sections of 5 tumor samples: P1, P3, P4, P5 and P7 where the classes of TIME in the tumor core were consistent with the corresponding pLCOs (Fig. 3, I to K, fig. S4, D and E). It is worth noting that the stroma showed very different immune cell infiltrating conditions. For example, condensed T cells appeared in the stroma of P3 (Fig. 3I) and P4 (fig. S4E) while the parenchyma was T cell excluded. Therefore, we developed an imaging processing method to quantify the fluorescent signals of the immune cells in the parenchyma regions (fig. S4G). The signals of T, Mph, and epithelial cells were all in good correlation with the average proportions of these cells in pLCOs derived from the same samples (Fig. 3L). We noticed that pLCOs derived from P4 can be divided into two groups according to the proportion of immune cells, i.e., the “desert” and the “cold” organoids. Consistent with this observation, the immunofluorescent staining of tissue section also revealed two types of architectures of the tumor parenchyma, the solid tumor with obvious macrophage infiltration and the lumenar structures with few immune cells (fig. S4, E and F). These results proved that the pLCOs well preserved the immune microenvironment of the parental tumor parenchyma and the diversities among the pLCOs reflected the spatial heterogeneity of the tumor parenchyma.

### Effector-like CD8^+^ T cells in pLCOs mediate the aPD-1 induced anti-tumor response

In contrast to scRNA-seq directly using dissected tumor tissues, in which only the patient response can be referred as the overall assessment for the anti-tumor performance of the immune cells, FascRNA-seq of single pLCOs allowed us to manipulate the treatment conditions and to evaluate the tumor killing functions of the small number of immune cells in individual organoids (Fig 4, A to D). Individual organoids showed diverse degrees of cell death, contributed by the drug treatment conditions as well as the heterogeneity in their immune microenvironments (Fig 4, B and C). Significant decrease in the survival index (Si: the ratio of living cells over total cells) were observed in P5 and P7 pLCOs (Fig. 4D), where the number and the proportion of ECs were remarkably reduced, suggesting effective anti-tumor immunity induced by aPD-1 (fig. S5, A to C).

To identify the immune cell types responding to aPD-1, we calculated the Pearson Correlation Coefficient (PCC) between the killing index (Ki = 1 – Si) and the cell proportion. At the patient level (i.e., pLCOs from the same patient were averaged), CD8^+^ T, CD4^+^ T, and NK cells all showed positive correlations with Ki under both aPD-1 and IgG treatments. Further analysis based on single pLCOs confirmed that only CD8^+^ T showed a positive correlation with Ki consistently across patients, while CD4^+^ T and NK behave differently in P5 and P7 pLCOs (Fig. 4E). Interestingly, NK showed a positive correlation under IgG but not aPD-1 treatment, consistent with the results of a previous report (*22*). Furthermore, the presence of CD8^+^ T cells significantly enhanced the Ki of pLCOs from all the samples, whereas CD4^+^ T and NK did not show such significant effect (fig. S5, D to F). Bulk study in conventional multi-well plates proved that anti-CD8 antibody (aCD8) eliminated the aPD-1 induced anti-tumor effect (fig. S5, G and H). Altogether, these data demonstrated that CD8^+^ Ts played the most important role in aPD-1 induced tumor immunity.

Next, we evaluated the functional status and response of these parenchyma infiltrating T cells (PITs) to aPD-1. Displaying of the CD8^+^ T cells in tSNE showed no distinguishable clusters associated with aPD-1 treatment or Ki of the parental organoids, whereas cells from different patients had minimum overlap (Fig. 4F), suggesting that the interpatient heterogeneity had a non-negligible impact on T cell transcription status and emphasizing the need for comparisons under the same patient background. Consistent with this observation, the enrichment of inhibitory and effector molecules was not seen (fig. S6A). We therefore referred to a recently published pan cancer T cell atlas (*23*) to identify the status of these T cells by comparing to the transcription signatures of the reported T cell subtypes. To our surprise, PCC analysis demonstrated that PITs were significantly similar to the C2 cluster of effector Ts featuring with the lack of inhibitory receptors and the upregulation of effector molecules, but not similar to the exhausted Ts (C1 and C7) (fig. S6, B and C).

Then, we analyzed the changes of transcriptome under the two treatment conditions (fig. S7A). Gene ontology (GO) analysis of the deferentially expressed genes (DEGs) revealed that pathways related to T cell activation, including the T cell receptor signaling and the PD-1 checkpoint pathways, were enriched under the aPD-1 treatment (fig. S7B). Consistently, genes related to T cell activation, including *OX40, TNFRSF9, CD96,* and *BCL2L1*, were significantly upregulated (Fig. 4G) and T cell proliferation was promoted (fig. S7, C and D). The expression of co-stimulatory receptors increased while the inhibitory receptors did not change (fig. S5, E and F). These results suggest that the aPD-1 treatment activates the PITs, promotes their proliferation, but has minus effect on their exhaustion status.

We investigated the effector molecules contributing to the tumor killing function of PITs. PCC between the average expression of the effector molecules in PITs and the killing index of their parental pLCOs were calculated. As shown in Fig 4H, the correlation with *NKG7* was most significant when treated with IgG, which explained why NK cells had a significant anti-tumor effect under this condition. In contrast, the top 3 effectors associated with aPD-1 were *GZMB, PRF1*, and *IFNG* (Fig 4, H to K), all of which were reported to play important roles in CD8^+^ T mediated tumor immunity (*24, 25*). The transcription level and the proportion of PITs expressing these three molecules were elevated with the aPD-1 treatment (fig. S7, G and H). Interestingly, the proportion of PITs co-expressing *GZMB* and *PRF1* increased for more than 2 times for both P5 and P7 pLCOs (Fig. 4I) and PITs expressing both molecules were correlated with higher level of Ki (Fig 4J), indicating the synergistic effect of the two molecules, consistent with previous reports (*26, 27*).

### Combining the phenotypic and transcriptomic data to identify the tumor-reactive T cells

Next, we tried to identify the tumor-reactive T cells which coexist with bystanders (*28*). We first calculated the average expression level of genes critical for CD8^+^ T activation and function for CD8^+^ Ts in individual pLCOs, and analyzed the PCC with Ki (fig. S6D). A total of 10 genes that had good correlation with Ki were chosen as a gene set to evaluate the functional status of CD8^+^ Ts (Fig. 5A), the average transcription levels of which were calculated as the T cell activation index (Tai). The average Tai in individual pLCOs was significantly correlated with Ki and upregulated upon the aPD-1 treatment (Fig. 5B, fig. S6E). Next, we calculated the Tai for all the CD8^+^ Ts and identified 29 cells with Tai > 0.5 (Fig. 5C). Comparison of the 29 cells with other Ts further confirmed their promoted activation level (fig. S6F). Consistently, pLCOs with the high Tai cells showed significantly elevated Ki (Fig 5D). Interestingly, the expression of inhibitory receptors was promoted in high Tai cells (Fig 5E), in agreement with previous reports on the more exhausted status of tumor-reactive T cells (*29–31*).

**Fig. 5.**
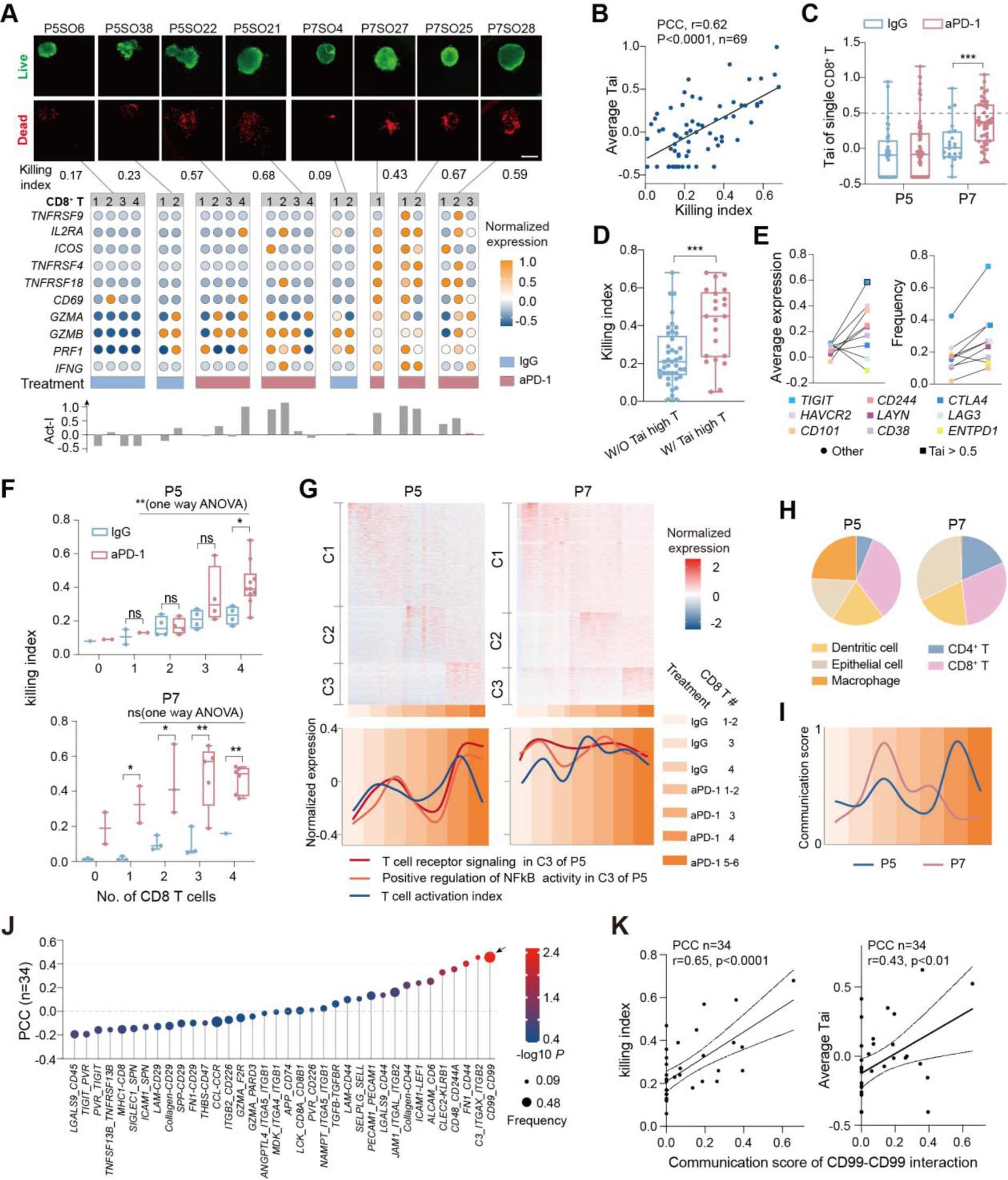
Identification of tumor-reactive T cells and cell-cell interactions promote T cell activation. **(A)** Images of single pLCOs stained with Calcein AM/PI (upper panels), bubble charts in the middle show the expression levels of the 10 genes for calculating Tai (each column represent a single T cell), bar graph at the bottom shows Tai of each T cells. Scale bars, 100 μm. **(B)** PCC analysis revealed significant correlation between average Tai and killing index for individual pLCOs. **(C)** Comparison of Tai for single CD8^+^ Ts under the two treatment conditions. **(D)** Comparison of killing index for pLCOs with (w/) or without (w/o) high Tai (>0.5) cells. **(E)** Comparison of the average expression levels (left) and frequencies (right) of the inhibitory receptors in high Tai cells and other CD8^+^ T cells. The black frame indicates gene with significantly different levels in the two cell groups. **(F)** Comparison of cell death under the two treatment conditions in pLCOs with different number of CD8^+^ T cells. **(G)** Upper panels: heatmap showing the dynamics of normalized gene expression in CD8^+^ T cells. Columns represent pLCOs ordered in treatment conditions and number of incorporated CD8^+^ Ts as indicated by the horizontal color bar at the bottom. All the genes are categorized into three groups. C1 includes genes upregulated in IgG treated pLCOs, C2 includes genes upregulated in aPD-1 treated pLCOs with 1-3 CD8^+^ Ts. C3 includes genes upregulated in aPD-1 treated pLCOs with 4 or more CD8^+^ Ts. Lower panels: average level of all the genes from the indicated GO terms for the indicated pLCO groups. Background color represents the pLCO groups indicated on the right. Line colors indicate GO terms or the Tai. **(H)** Pie chart showing the abundance of cell-cell interactions received by CD8^+^ Ts and sent by different types of cells. **(I)** Average communication scores of CD8^+^ T-CD8^+^ T interactions for the indicated pLCO groups. Background color represents the same pLCO groups as in panel **g**. **(J)** Analysis of PCC between the communication scores of ligand-receptor pairs and killing index of P5 pLCOs. Note the homophilic interaction of CD99 was the most significantly correlated with killing index. **(K)** Correlation between CD99 homophilic interaction and killing index (left) or average Tai (right) of P5 pLCOs. The center lines in the box plots of **(C, D and F)** represents the median values. The bounds of box represent the median values of the upper half and the lower half. The bounds of whiskers represent the maxima and the minima. P values were determined by unpaired, two-sided student’s *t* tests unless otherwise indicated, *P < 0.05, **P < 0.01, ***P < 0.001.

### Synergies of CD8^+^ T cells enhanced the immune activating effect of aPD-1

We observed that in P5 pLCOs, the significant response to aPD-1 can only be seen in organoids with 4 or more CD8^+^ Ts, whereas in P7 pLCOs, a single CD8^+^ T is sufficient to produce an obvious cytotoxicity effect (Fig. 5, A and F). We then analyzed whether there are gene expression programs regulated by the number of T cells. All the genes were divided into three groups: upregulated in IgG treated control organoids (C1), aPD-1 treated organoids with 1-3 CD8^+^ Ts (C2) and 4 or more Ts (C3). C3 of P5 was enriched with genes corelating to T cell activation (GO terms “T cell receptor signaling” and “positive regulation of NF-kB transcription factor activity”) while C3 of P7 was not (tables S1), suggesting promoted T cell activation with increasing number in P5 organoids. To further illustrate the dynamics, the average expression level of genes within these GO terms was plotted over CD8^+^ T number in single organoid. For P5 organoids, CD8^+^ T activation was promoted in pLCOs with more T cells in both the control and the aPD-1 treatment conditions while such dynamic was not observed in P7 organoids (Fig. 5G). Consistently, Tai also increased with T cell number in P5 but not P7 organoids. These results suggest a synergistic effect between T cells which promote their activation and sensitivity to aPD-1. Yet it seems that this synergistic effect had an impact only when T cell activation was insufficient, since this phenomenon was not observed in P7 organoids where T cells were activated more thoroughly (Fig. 5, A, F, and G).

To further corroborate whether the synergistic effect between CD8^+^ Ts has a molecular base, cell-cell interactions in individual organoids received by CD8^+^ Ts were elucidated through bioinformatic analysis (fig. S8, A to D). Consistent with the synergistic effect mentioned above, interactions between CD8^+^ Ts were most abundant in P5 pLCOs (Fig. 5H) and the communication score of CD8^+^ T-CD8^+^ T interactions had similar dynamics as Tai (Fig 5, H and I), while the MHCI-CD8 interactions between DC and CD8^+^ T were more abundant in P7 pLCOs, which may explain the more sufficient T cell activation (fig. S8, A, B, and E). Next, we tried to figure out the possible ligand-receptor pairs that might contribute to the synergistic effect of CD8^+^ Ts. The PCC analysis in P5 pLCOs revealed that the homophilic interaction of CD99 was the most significantly correlated with Ki and Tai (Fig. 5, J and K). CD99 is a transmembrane protein broadly expressed in many cell types(*32*). Evidences on its costimulatory role in T cell activation have been reported (*33, 34*). Stimulation of CD99 with agonistic antibodies enhanced the expression of several T cell activation markers and sensitizes cells for activation through the CD3 signaling (*35*). Consistent with these reports, P5 pLCOs with T cells receiving CD99 homophilic interactions showed significantly enhanced Ki (fig. S8F). In contrast, the effect of CD99 was not so significant in P7 pLCOs (fig. S8G). These data suggested that the homophylic interaction of CD99 might play a role in T cell activation in a suboptimal immune microenvironment.

### Gathering of M2-like macrophages prevents the accumulation of T cells

pLCOs derived from patient samples vary in their immune cell components, yet such variations were not completely random but exhibited some general and patient specific characteristics. Among the pLCOs with immune cells, the high frequency of myeloid cells, mostly Mphs, was associated with inhibited lymphocyte accumulation and compromised response to ICB (Fig. 6, A and B). The “repellency” of macrophages to Ts was also observed on tissue sections, as quantification of the macrophage (CD163) and the T cell signals (CD3) in the parenchyma region demonstrated the negative correlation between the two cell types (Fig. 6, C to E).

**Fig. 6.**
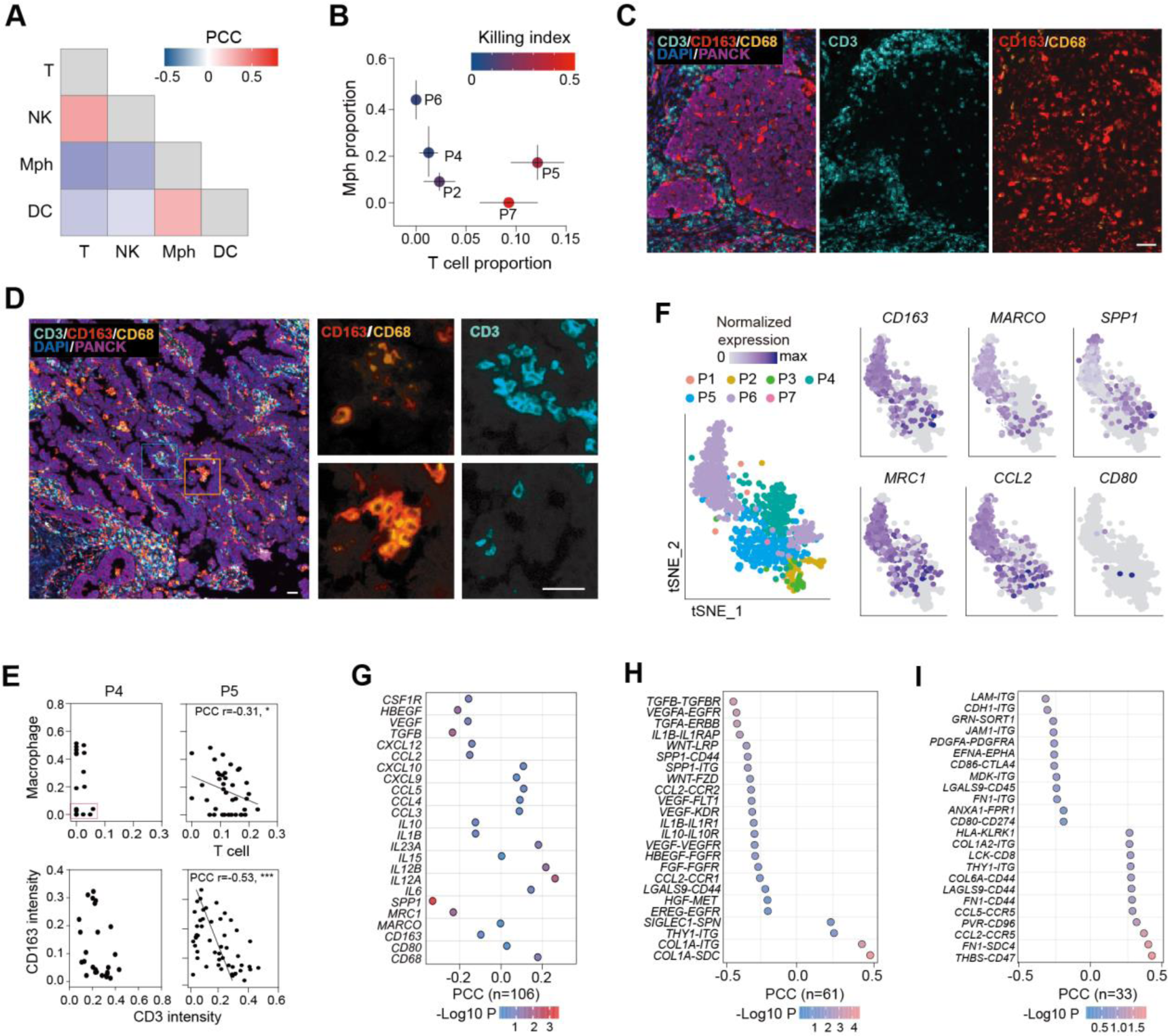
Gathering of M2-like macrophages prevents the accumulation of CD8^+^ T cells. **(A)** Heatmap showing PCC between the proportions of different immune cell types. Note the negative correlation between Mph and T cells. **(B)** Dot plots showing the average proportions of T and Mph in the pLCOs from different patient samples. The lines on the dots represent the standard deviation and the color of the dots indicates the average Ki for pLCOs from one sample. **(C and D)** IHC images of P4 **(C)** and P5 **(D)** tumor tissue sections. Note the excluding of T cells from tumor parenchyma of P4 and the presence of both T and Mph in P5. Scale bars: 60 μm. **(E)** Quantification of T cell and Mph proportions in individual pLCOs (upper panels) and the CD3 and CD163 signals in randomly picked 0.2mm x 0.2mm regions of the parental tumor parenchyma (lower panels). Note the similar trends of the T over Mph ratio in individual pLCOs and randomly picked parenchyma regions of the same patient sample. **(F)** The tSNE projection of Mph from all the pLCOs. Data points of each cell were labeled with patient or normalized expression of indicated genes. **(G)** PCC Between T cell number and average expression level of immune related genes in the Mphs in individual organoids. **(H and I)** The ligand-receptor pairs between Mph and ECs **(H)** or Mph and T cells **(I)** that had the most significant correlation with the number of T cells in individual pLCOs. Note the negative correlation for most of the ligand-receptor pairs between Mph and ECs.

Displaying the transcriptome data of macrophages in tSNE revealed obvious M2 signatures (Fig 6F), including *CD163, MRC1,* and *MARCO*, while the M1 signatures, *CD80, CD86,* and *CD68,* were detected in very low frequencies (Fig. 6F). Consistent with the transcriptome data, immunofluorescent staining of the tissue sections also demonstrated M2-like tumor-associated macrophages (TAMs) as the majority (CD163+ TAMs dominant). Furthermore, TAMs in P4 showed more M2-like polarity as indicated by the ratio of CD163 over CD68 signals, in agreement with the more complete T cell elimination (Fig 6, C to E). PCC analysis revealed that the expression level of M2 signature genes, EMT and angiogenesis-promoting genes (*VEGF, TGFB*), immune suppressive cytokines *(IL10, IL1B)*, and monocyte recruitment chemokines (*CCL2, CXCL12*) were negatively correlated with T cell accumulation, while the proinflammatory cytokines, such as *IL12*, and the chemokines promoting T cell recruitment had a positive correlation with T cell accumulation (Fig. 6G). Cell-cell interaction analysis revealed that these molecules compromise T cell infiltration largely through the tissue remodeling associated interactions between Mphs and ECs (Fig.6H), while interactions between Mph and Ts are two-faced, exerting both positive and negative effects (Fig. 6I, fig. S9A).

Lastly, we displayed the cell-cell interaction networks in Sankey diagram to show the local TIME in the tumor parenchyma for individual patients. The interaction networks were more immune suppressive in P6 and P4 compared to P5 (fig. S9, B to D). For P5, macrophages had mixed characteristics of M1 and M2 polarity. Correspondingly, the cell-cell interaction networks in individual organoids vary from immune suppression to immune activation. For example, in the PD-1-sensitive organoid P5SO22, MHCI-CD8 mediated antigen presentation and CD86-CD28 mediated co-stimulation exist (fig. S9E). Interestingly, we noticed that although the cell-cell interaction networks in P6 were mostly immune inhibitory, interactions between Mphs and DCs were favorable for DC maturation, including *CD40LG-CD40* interactions. In addition, these interactions enhanced significantly upon the aPD-1 treatment (fig. S9D). Accordingly, the transcription level of genes related to DC maturation, including *MHCI, MHCII, CD80/86, CD40*, and *CCR7*, elevated significantly and the proportion of DCs decreased upon the aPD-1 treatment (fig. S9, F and G).

## Discussion

In this study, we developed a single cell processing and RNA sequencing platform (FascRNA-seq) featured with 1) automatic single cell/organoid processing and distribution with a minimal cell loss and a minimal starting cell number, and 2) the capability of tracing single cells back to their parental organoids, simply by recording the microwells where the cells and the barcode sequence delivered. FascRNA-seq is suitable for analyzing samples containing no more than 1000 of cells, for example individual organoid, especially when there are a small number of pivotal cells, such as functional immune cells or stem cells. Here, we evaluated the response of individual tumor organoids to ICB, at the meanwhile, captured the immune cells responsible for these changes, and assessed their transcription profiles. In addition, our methods can also be used for the single cell analysis of normal organoids developed from stem cells (*36*).

Unlike organoids developed from single “seed” cells, for example Lgr5^+^ intestine stem cells (*37*), which are composed of pure epithelia, our pLCOs are composed of tumor cells and the tumor infiltrating immune cells, owing to the digestion free sample processing method, by which clusters of tumor cells are isolated directly and the immune infiltrates are retained. Tumor is a heterogeneous tissue composed of parenchyma and stroma, both of which have a huge diversity in their immune infiltrates (*2, 38*). We demonstrated that the average proportions of immune cells in the pLCOs are in good agreement with the overall level of immune infiltrates in the parenchyma, but different from that in the whole tumor tissue containing abundant stroma. In addition, we also proved that the differences among individual pLCOs derived from the same sample recapitulate the intra-parenchyma heterogeneity, for example, the “desert” and “cold” pLCOs derived from P4, representing the two types of parenchyma architecture. Therefore, we conclude that pLCOs retain the local TIME in tumor parenchyma where it is derived.

As an in vitro model, pLCOs provide two advantages. First, multiple pLCOs can be derived from a tumor sample, providing *in vitro* models with the exact same patient background (i.e., recapitulating the fundamental features of parental tumor, such as the genetic mutations and the classes of immune infiltrates) but with slight variations in TIME (i.e., corresponding to the heterogeneity of local microenvironment in the tumor core). Such diversities will help us define the patient-specific factors affecting the anti-tumor immune response, such as the synergistic effect of CD8^+^ Ts we observed in P5, which may otherwise be overwhelmed by the huge difference among patients. Second, the operability and traceability of pLCOs render the convenience to investigate immune cell dynamics under different treatment conditions, which can help us discover next-generation immunotherapies and understand the mechanism of action more efficiently.

Previous studies using scRNA-seq unraveled the heterogeneity of tumor infiltrating lymphocytes (TILs) (*39–42*), and the presence of exhausted T cells featuring with loss of effector molecules and co-expression of multiple inhibitory receptors (*42, 43*). Yet, the spatial distribution of T cells with different status are unclear. Here we observed that the parenchyma infiltrating CD8^+^ Ts are effector-like rather than exhausted. Our other study on breast cancer also demonstrated that compared to those isolated from fresh tumor tissues, CD8^+^ Ts in the primary tumor organoids are less exhausted. One possible explanation to this observation is that the distribution of exhausted T cells is uneven between tumor parenchyma and stroma. In the future, it will be interesting to analyze immune cells from other parts (i.e., the stroma and the tertiary lymphoid structures) by modifying the tissue processing method to splice the immune landscape of the whole tumor tissue, and furthermore, to elucidate what subtype of CD8^+^ Ts can infiltrate into the parenchyma and what kind of molecular features they have.

The current pLCO model represents the local TIME in tumor parenchyma, but lacks the surrounding stroma which contains abundant immune components, such as the tertiary lymphoid structures and blood vessels. In the future, the pLCO model can be further improved by co-culturing, for example, with the peripheral blood or tumor infiltrating lymphocytes and by constructing more sophisticate 3D structures representing the stromal tissue. Nevertheless, using the pLCO model alone coupled with FascRNA-seq we are able to 1) elucidate change of TIME induced by the anti-PD-1 antibody, 2) identify tumor-reactive T cells through the evaluation of a set of biomarker genes in combination with the phenotypic data of the parental organoids, 3) unravel the factors regulating T cell activity, including a mutual activation effect between CD8^+^ Ts which is probably mediated by the CD99 homophilic interactions.

## Materials and Methods

### Design, fabrication, and operation of MoSMAR-chip system

The modular superhydrophobic microwell array chip (MoSMAR-chip) (fig. S1A) was fabricated using standard injection-molding method with polystyrene (Hochuen Technologies, Shenzhen, China). Plasma cleaning was performed to make the surfaces of the entire MoSMAR-chip hydrophilic, suitable for cell culture. Next, a home-made superhydrophobic paint (materials were listed in tables S2) was prepared as follows: 1 g of 1H, 1H, 2H, 2H-perfluorooctyltriethoxysilane (Sigma-Aldrich) was first added into 99 g of absolute ethanol and mechanically stirred for 2 h. Then, 6 g of titanium oxide nanoparticles (TiO2, ∼60 to 200 nm) (Sigma-Aldrich) and 6 g of P25 TiO2 (∼21 nm) (Degussa) were added into the solution to make a paint-like suspension, which was sonicated for 30 s to disperse the particles. After that, the suspension was pipetted onto the recessed top surfaces of the reaction chip and the transfer coverslip of the MoSMAR-chip and air-dried completely within 30 s. Finally, the MoSMAR-chip was autoclaved and sealed in a plastic bag until use.

The operation of the MoSMAR-chip is flexible. First, the number of the reaction chips assembled into the chip frame can be adjusted according to the throughput demand. Second, the loading of nl-scale reagents into the transfer coverslip can be achieved using a non-contact acoustic liquid handling system Echo 550 (Beckman Coulter) (fig. S1C). Third, the loading of reagents into the microwells of the reaction chip can be carried out either as a whole using the immerse-aspirate method or individually using the spot-cover method with the transfer coverslip, which contains pre-stored reagents (fig. S1D). Finally, as described previously, cells can be loaded into the microwells by either random seeding or droplet rolling, and cultured in either the immersion mode or the droplet mode.

### Construction and optimization of the automated single-cell distribution instrument

The home-built single-cell distribution instrument (SCDI) consisting of an integrated microscopy module for imaging cells, a six-axis motion and control module for operating the chip platform and the robotic arm, and a motor-controlled hydraulic microneedle module for cell delivery. The microscopy module consists of an objective lens (Olympus), a digital camera (MER-U3-L, Daheng Optics), and a series of optical path conversion parts (listed in the tables S3) to transfer images from the objective to the camera. The digital camera is linked to a PC-based graphics card (GeForce RTX 3080, NVIDIA) for imaging processing. The motion and control module includes a three-axis motion platform for chip positioning and a three-axis robotic arm for controlling the capillary movement, each of which is constructed using three 57 three-phase stepper motors (Daheng Optics, GCD-202150M). The hydraulic module is constructed with a microinjector (CellTram vario, Eppendorf) motorized by a 57 three-phase stepper motor. A glass capillary electrode (B100-75-10, Sutter) pulled by a micropipette puller (P-1000, Sutter) is installed onto the microinjector for extracting and releasing single cells or organoids. The synchronized control of all the movements is achieved through the use of a stepper motor controller (Daheng Optics, GCD-040102M) that communicates with the control program via a CAN (Controller Area Network) bus.

The core of the custom-made software is a real-time target recognizer based on the YOLOv4 neural network framework, implemented in C++ language using the OpenCV computer vision library (fig. S2A). To construct the training set, we manually captured 600 various single-cell images using the single-cell distribution instrument and label the regions of interest (ROIs), including the capillary microneedle tips and single cells, using the labelImg software. The training set was fed into the YOLO neural network for iterative training of the recognizer. The training process was considered complete when the loss function value fell below 0.6. The trained and the neural network code can be obtained from the Github.

The system optimization of the SCDI is focused on: the recognition accuracy of the neural network recognizer, the size of the neural network training set, the Ficoll ratio in cell samples, the cell density, the precision of the hydraulic-driven single-step movement, and the inner diameter of the capillary microneedle. The optimized parameters were determined as follows: the neural network was trained with a dataset of 600 samples; the capillary micro-needle with an inner diameter of approximately 100 μm was driven with a single-step precision of 50 nl and the cell solution samples were prepared with a concentration of 1×10^5^ cells/ml in 5% Ficoll solution.

### Single-cell transcriptome library construction

The single-cell RNA-seq protocol of the FascRNA-seq is modified from the Grouped-seq developed previously by our group. All the reagents used in the protocol are listed in the tables S4. Briefly, the 114-base-long capture oligo that attaches to the streptavidin-coated Dynabeads^TM^ is constructed by linking two short oligos (P1 and P2) together using a linker via enzymatic ligation (Fig.2A). We constructed 192 capture oligos with different barcodes. We first employed an Echo 550 instrument to load the beads and the capture oligos into the microwells of the transfer coverslips of the MoSMAR-chip. In each microwell, 500 nl of the oligos with a concentration of 10 ng/mL were reacted with 80 nl of M270 magnetic beads, diluted at a 1:2 ratio. The loaded transfer coverslip was then stored at 4°C until use.

Following the single-cell loading with the single-cell distribution instrument, the transfer coverslip containing the magnetic beads and the barcoded capture oligos was aligned and covered onto the reaction chip upside down. After the loading of the barcoded-beads was finished, the transfer coverslip was removed. Next, the reaction chip was washed with 5x RT buffer to remove any unlinked capture oligos from the microwells. Then, 200 μL of lysis buffer (200 mM Tris–HCl, 20 mM EDTA, 1% Sarkosyl, 50 mM DTT) was pipetted to cover the microwell array region of the reaction chip with. The cells were lysed *in situ* by the diffusion of the lysis buffer into the microwells. The magnetic beads, which were in close proximity to the cells, immediately captured the released mRNAs. Following a 15-minute incubation, the beads were gently washed off from the MoSMAR-chip with 1 ml of 6× SSC solution and collected into a 1.5-ml centrifuge tube. Subsequently, the beads were subjected to two washes with 600 μl of washing buffer A (10 mM Tris–HCl, 15 M NaCl, 1 mM EDTA, and 0.1% Sarkosyl), one wash with 300 μl of washing buffer B (10 mM Tris–HCl, 15 M NaCl, and 1 mM EDTA), and one wash with 50 μl of 5× Maxima H RT buffer (Thermo Fisher). Finally, the washed beads were resuspended in 4 μl of 5× Maxima H RT buffer and kept on ice for further processing.

The magnetic beads were loaded into 16 μl of reverse transcription (RT) mix, comprising 10.5 μl of Nuclease-Free water, 2 μl of 10× dNTP, 2 μl of template switch oligo (TSO, 25 μM), 1 μl of Reverse Transcriptase (200 U/μl), and 0.5 μl of RNase inhibitor (40 U/μl) (Thermo Fisher). Subsequently, the mixture was incubated at room temperature for 30 min, followed by 42°C for 90 min to generate cDNA. Alternatively, if desired, ten more extension cycles (50°C for 3 min, 42°C for 3 min) could be performed to improve cDNA yield. The beads were then subjected to one wash with 20 μl of TE/SDS (1× TE and 0.5% sodium dodecyl sulfate), two washes with 20 μl of TE/TW (1× TE and 0.01% Tween 20), one wash with 20 μl of ddH_2_O, and finally resuspended in 12 μl of ddH_2_O.

The beads were supplemented with a PCR mix comprising 20 μl of 2× HotStart Readymix (Kapa Biosystems), 4 μl of 5′ end biotin-modified P7 primer (10 μM, tables S5), and 4 μl of TSO primer (10 μM, tables S5). The PCR program consisted of an initial denaturation step at 95 °C for 3 min, followed by 4 cycles of 98 °C for 20 s, 65 °C for 45 s, and 72 °C for 3 min, and subsequently, 23 cycles of 98 °C for 20 s, 67 °C for 20 s, and 72 °C for 3 min. A final extension step was performed at 72 °C for 5 min. The PCR products were purified using 0.6× AMPure XP beads (Beckman Coulter) and eluted into 50 μl of ddH2O.

PCR products were fragmented using the Covaris M220 system (Covaris) to generate 3′-end cDNA fragments. Then, end repair, 5′ phosphorylation, dA-tailing, adaptor ligation, size selection, and PCR enrichment were performed sequentially using the NEBNext Ultra™ II DNA Library Prep Kit for Illumina (NEB). Lastly, PCR products were purified using the AMPure XP system and the quality of the library was assessed using the Agilent Bioanalyzer 4200. The libraries were subjected to paired-end sequencing (150 nt each) on the HiSeq-PE150 instrument (Illumina). Two paired-end reads, Read 1 containing a sequence typically mapping to the 3′ end of an mRNA transcript and Read 2 containing a wellcode (16 bases) identifying a specific microwell along with a UMI (10 bases) were obtained. A single MoSMAR-chip can sequence up to 768 single cells. The primers and barcode sequences designed for FascRNA-seq were listed in tables S5.

### Generation, culture, drug treatment, and single-cell digestion of patient-derived tumor organoids

The lung cancer samples were obtained from the Peking University People’s Hospital with approval from the local Ethics Committee. Main inclusion criteria included patients with clinically local advanced or metastatic lung cancer, aged 18 years or older, fresh tissues available through surgical resection of the lesions. Candidates were assessed to determine eligibility and informed consents were obtained before operation. Upon collection, the lung cancer tissues, which had been stored in a preservation solution (Dulbecco’s modified Eagle’s medium/Nutrient Mixture F12 (DMEM/F12) containing 1% penicillin and streptomycin (Gibco)), were transported to the laboratory and processed within 24 hours. Initially, the lung cancer tissues were documented, weighed, and measured for volume. Then, the tissues were sectioned into multiple 1– 5 mm^3^ pieces without bias. The pieces were minced into small fragments using surgical scissors, followed by resuspension in 10 ml of preservation solution. The suspension was filtered through a 100-μm strainer (Falcon) with gentle grinding and pushing of tissue masses through the strainer using a 5-ml syringe plunger. After rinsing, any residual impurities on the membrane were discarded along with the strainer. Subsequently, the filtrate containing cell clusters and single cells was strained through a 40-μm strainer to collect the cell clusters with sizes ranging from 40 to 100 μm. The filter membrane was separated from the strainer, placed in 2 ml of LCOM (Lung cancer organoid medium), and thoroughly washed with a pipette to release the cell clusters into the media. The membrane was discarded, and the cells were cultured in suspension overnight.

For culturing pLCOs in a multi-well plate, the pLCOs in suspension were centrifuged at 500 × g for 5 minutes at 4 °C and then resuspended in cold growth factor-reduced Matrigel (BD Biosciences). Subsequently, 60-μl drops of the Matrigel cell cluster suspension were plated onto ultra-low attachment 96- or 48-well plates with a flat bottom (Corning) and allowed to solidify at 37 °C for 20 minutes. The seeding density was adjusted to approximately 500 organoids per well. Once the Matrigel had solidified, 100 μl of LCOM was added to each well, and the plate was transferred to a cell culture incubator at 37 °C with 5% CO2. The LCOM medium was refreshed twice a week. The composition of the LCOM medium can be found in tables S6.

Individual tumor organoids were inoculated into the microwells of the MoSMAR-chip using the single-cell distribution instrument. Next, the tumor organoids were treated with the immune checkpoint inhibitor Nivolumab (Selleckchem, Cat#A2002) and the Ultra-LEAF Purified Human IgG4 Isotype Control Recombinant Antibody (Biolegend, Cat#403701) at a concentration of 10 μg/mL. After a culture period of 5-7 days, the tumor organoids were moved one by one from the microwells to the transfer wells, where the organoids were digested into single cells by a 20-min incubation at 37°C with TrypLE solution. Then, gentle pipetting with a 20-μl pipette tip was performed for about 60 times to completely digest the tumor organoids. After a settling period of 10 minutes, the single-cell distribution instrument was employed to distribute these cells into microwells to form a single-cell array for subsequent RNA sequencing. All of the tumor organoids treatment related reagents can be found in tables S7.

### Immunohistochemical staining of patient’s tumor tissue slices and immunofluorescence staining of tumor organoids in frozen sections

Immunohistochemistry staining was performed on paraffin slides of lung tissue with 2 μm thicknesses. The slides were initially heated at 72 °C for 30 minutes to ensure proper tissue adhesion.

The slides were deparaffinized in xylene for 10 mins and repeat three times, and rehydrated in absolute ethyl alcohol for 5 mins and repeat twice, 95% ethyl alcohol for 5 mins, 75% ethyl alcohol for 2 mins, sequentially. Then the slides were washed with distilled water 3 times. A microwave-oven is used for heat-induced epitope retrieval. During epitope retrieval, the slides were immersed in boiling EDTA buffer for 15mins. Antibody diluent and block buffer (AlphaX Biotech) was used for blocking. The mIHC staining part was performed and analyzed according to a 6-plex-7-color panel. All the primary antibodies (CD3, CD68, PANCK, PDL1, CD163, and CD56) were incubated for 1 hour at 37°C. Then slides were incubated with Ploymer HRP (AlphaX Biotech) for 10 mins at 37°C. AlphaTSA Multiplex IHC Kit was used for visualization. After each cycle of staining, heat-induced epitope retrieval was performed to remove all the antibodies including primary antibodies and secondary antibodies. The slides were counter-stained with DAPI for 5 mins and enclosed in Antifade Mounting Medium (NobleRyder). Axioscan7 (ZEISS) was used for imaging the visual capturing. Detailed information of the antibodies can be found in tables S7.

The cell proportions of tumor tissues were evaluated based on the immunohistochemistry staining images with ImageJ software. Specifically, the tumor parenchyma as image ROI was inferred according to the merged image channel, then the ROI was applied to the channel of DAPI signal and excluded the stroma region outside ROI. The threshold was applied to acquire cell nucleus signal of tumor parenchyma as base value, followed by quantification of various cell types including T cells, NK cells, epithelial cells, and macrophages in the same way.

Tumor organoids were harvested from Matrigel by centrifugation at 800 rpm once they reached an approximate diameter of 100 μm. Subsequently, they were embedded in a 10 mg/mL Fibrin matrix and fixed in a 4% paraformaldehyde solution for 30 minutes at room temperature. To facilitate cryopreservation, the fixed organoids were dehydrated overnight at 4°C in 30% sucrose. Following the removal of sucrose, the tumor organoids were carefully embedded in cryomolds using optical coherence tomography (OCT) compound and rapidly frozen at −20°C. Cryosections with a thickness of 25 μm were obtained and affixed onto charged slides. To ensure optimal immunostaining, the sections were permeabilized and blocked in TB buffer (TBST + 0.5% TritonX-100 + 1% BSA) for 1 hour at room temperature. Primary antibody staining was performed by incubating the slides overnight at 4°C with specific antibodies, including EPCAM (Abcam) and CD3 (Abcam). Following primary antibody incubation, the slides underwent further staining with secondary antibodies, including Alexa Fluor 488-conjugated anti-mouse IgG (Invitrogen), Alexa Fluor 647-conjugated anti-rabbit IgG (Invitrogen), and DAPI (Invitrogen) for 1.5 hours. To visualize and analyze the stained samples, fluorescence images were obtained using a Nikon A1HD25 microscope. Detailed information of the antibodies can be found in tables S7.

### Quantitative evaluation of drug sensitivity response of pLCOs and cell proportion of tumor tissue

The drug sensitivity response of tumor organoid models was evaluated based on the cell viability, which was measured using Calcein AM/PI staining reagents. Briefly, 1 μl of AM solution and 3 μl of PI solution were added to 100 μl of staining buffer. The mixture was then gently oscillated in dark for approximately 1 minute using an oscillator, followed by the dilution with PBS in a 1:30 ratio. Subsequently, the staining solution was dispensed into the microwell array of the transfer coverslip with a volume of 500 nl per well using the Echo 550 acoustic liquid handling system. Next, the MoSMAR-chip containing the tumor organoids was gently immersed in approximately 2 ml of PBS solution and then the liquid was aspirated, resulting in the washing and replacement of the reagent in all the microwells. After that, the transfer coverslip containing the staining solution was covered onto the reaction chip, followed by incubation in a cell culture incubator for approximately 15 minutes in a light-protected environment. The reaction chips were then retrieved and subjected to bright-field, green, and red fluorescent imaging using an inverted fluorescence microscope (IX83, Olympus). The populations of live and dead cells were quantified by measuring the total cell area for each dye with ImageJ software. Meanwhile, the area of green signal was considered as the overall size of each tumor organoid. The response level of the tumor organoids was determined by subtracting the quantified signal of all dead cells from the quantified signal value of all live cells, which was then divided by the overall size of the tumor organoid.

### Flow cytometry analysis for tumor organoids

To assess the abundance of T cells within the samples, tumor organoids were dissociated using TrypLE Express enzyme. To minimize non-specific binding, Fc receptors (FcR) were blocked by incubating the cells with Human BD Fc Block (BD Biosciences, 1:100) for 10 minutes in FACS buffer (PBS + 2% FBS). The dissociated cells were then washed with FACS buffer and stained with specific antibodies at 4°C for 30 minutes. The antibody panel included Pacific Blue™ anti-human CD45 Antibody (BioLegend), BV510 Mouse Anti-Human CD3 (BD Biosciences), PerCP-Cy™5.5 Mouse Anti-Human CD8 (BD Biosciences), and FITC Mouse Anti-Human CD4 (BD Biosciences). Following the staining procedure, the cells were washed twice with FACS buffer, and a near-infrared viability dye (Invitrogen) was added to exclude dead cells. The cell suspensions were then analyzed using the Aria III flow cytometer (BD Biosciences). Compensation beads stained with corresponding single antibodies were used to establish compensation settings. The acquired data were subsequently analyzed using FlowJo v.10.8.1 software, and only single, viable cells were included in the analysis. This flow cytometry-based approach allowed for the quantification and characterization of T cells within the dissociated tumor organoids. By examining the expression of CD45, CD3, CD8, and CD4, the population of T cells and their subtypes could be identified and quantified. Detailed information of the antibodies can be found in tables S7.

### Single-cell RNA-seq analysis of tumor organoid models

The bioinformatics analysis pipeline can be found on Github. Read 1 Fastq files were aligned to an appropriate reference genome using STAR v2.4.0a with default settings. For mouse cells, the mm10 reference genome was employed, while for human cells, the hg38 reference genome was utilized. In the cases involving mixtures of mouse and human cells, the union of hg38 and mm10 references was used. Read 2 Fastq files were used to extract wellcodes and UMIs based on the wellcode-UMI-Poly T pattern (16-nt wellcode, 10-nt UMI, 16-nt poly T). The RNNS algorithm was developed for wellcode identification, which allowed either perfect match or a 1-nt mismatch with the reference. Since the designed wellcodes were at least 2-Hamming distance apart from each other, a 1-Hamming distance difference between the designed and measured wellcodes was permitted to correct errors arising from PCR or sequencing. PCR duplicates were identified when the same wellcode, UMI, and gene ID (ENSEMBL human GRCh38, mouse GRCm38) were detected. The gene abundances of the samples were estimated using featureCounts (Version 1.5.0-p1).

For the integrated dataset comprising all seven primary single tumor organoid models derived from clinical sources, all individual single-cell samples with >300 transcripts and <20% mitochondrial reads were retained in the analysis. After the aforementioned quality control steps, the filtered single-cell transcriptomic expression matrix obtained from the data was subjected to further analysis using Seurat (*44*) (version 4.2.0). The Seurat object was normalized for the identification of variable genes by running NormalizeData and FindVariableFeatures (SeuratObject, selection. method = “vst,” features = 1500). Next, the harmony algorithm (*45*) was applied to integrate samples based on patients and to therefore eliminate batch effects. The Seurat object holding integrated batch effect corrected expression matrix was returned. The integrated dataset was scaled and the principal components (PCs) was computed using default settings. UMAP dimensional reduction via RunUMAP() and FindNeighbors() was performed using the first 10 principal component analysis (PCA) dimensions as input features. FindClusters() was computed at a resolution of 0.8. Additionally, the tumor organoid line dataset was calculated according to the same pipeline and the corresponding pseudotime analysis was performed using Monocle 3 (*46*) (version 1.2.9). The differential gene expression analysis based on metadata was identified using FindAllMarkers() using default parameters. Furthermore, the cluster-specific pathways were revealed by Kyoto Encyclopedia of Genes and Genomes (KEGG) pathway enrichment analysis and Gene Ontology (GO) enrichment analysis for biological processes completed by the clusterProfiler (*47*) (version 4.4.4).

Cell type annotations were performed through SingleR (*48*) (version 1.10.0), which assigns cellular identity for single cell transcriptomes by comparison to reference datasets. For the single-cell samples in the dataset, their cell types were initially identified using two datasets, BlueprintEncode and HumanPrimaryCellAtlas, included in SingleR. The identified immune cells were further validated using five additional datasets, including ImmuneCellExpressionData, Hematopoietic, Monaco, MonacoImmuneData, and Novershtern HematopoieticData. In this workflow, the integrated cell type labels were further refined by combining with the expression features of Seurat clusters, ultimately achieving the identification of cell types across the entire dataset. Based on this workflow, major cell types (Epithelial cells, Macrophages, Endothelial cells, Monocytes, Dendritic cells, NK cells. CD8^+^ T cells, CD4^+^ T cells, Fibroblasts, B cells, etc) were identified.

For the non-immune cell types identified in the above dataset, we used inferCNV (*49*) (version 1.12.0) to estimate the chromosomal CNVs following the standard workflow for 10× genomics with default parameters. Due to the relatively homogeneous origin of our tumor organoid models, we set ref_group_name=NULL in order to make the average expression of all non-immune cells in the dataset as a reference. Then, the inferCNV results of each cell were quantified as a CNV score. Specifically, the complete loss copy or addition of more than one copy were counted as two points, the loss of one copy or addition of one copy were counted as one point, and the neutral situation was not counted in the CNV score.

The ligand-receptor interaction based cell communication analysis of each single-organoid level sub-database was completed by two R packages, CellCall (*50*) (version 0.0.0.9000) and CellChat (*51*) (version 1.6.1). The CellChat algorithm was mainly applied to ECM-receptor and cell-cell contact two aspects of inference and calculation, the CellCall algorithm was mainly used for secreted signaling related ligand-receptor calculation and enrichment of transcript factor based on the expression of cell receptor related genes. Specifically, in CellChat package related analysis, we sequentially applied identifyOverExpressedGenes and identifyOverExpressedInteractions with lenient default parameters to find the ligand-receptor gene combinations overexpressed in major cell types. Next, we ran computeCommunProb (with nboot = 1000) followed by computeCommunProbPathway, netAnalysis_computeCentrality, and aggregateNet with default parameters to find the ligand-receptor pathways present. Lastly, we merged the case and control objects and ran computeNetSimilarityPairwise with type “functional”. In CellChat package analysis, the sub-datasets of single-organoid were firstly generated as Seurat object, then the CreateObject_fromSeurat and TransCommuProfile procedure were performed with default parameters successively in order to infers intercellular communication by combining the expression of ligands/receptors and downstream TF activities for certain L-R pairs. The calculated single-organoid based ligand-receptor interaction results of CellChat and CellCall were derived in order to apply further aggregated and statistically analysis.

### Statistical analyses

Statistical analyses, including significance analysis and correlation analysis, were conducted using Microsoft Excel or GraphPad Prism 8 software. The data are presented as the means ± SD or the means ± SEM, and P values were calculated. For additional statistical significance values and sample sizes in graphs, refer to the figure legends and Materials and Methods for more details.

