## Supplementary materials for "Function-associated scRNA-seq on single lung cancer organoids unravels the immune landscape of tumor parenchyma"

Chang Liu *et al.*

Xiaofang Chen,
Jun Wang,

**This PDF file includes:**

Figs. S1 to S9
Legends for Figs. S1 to S9
Legends for tables S1 to S7
Description for Movies S1 to S3

**Other Supplementary Materials for this manuscript include the following:**
Supplementary tables S1 to S7
Movies S1 to S3

23 ## Supplementary Figures

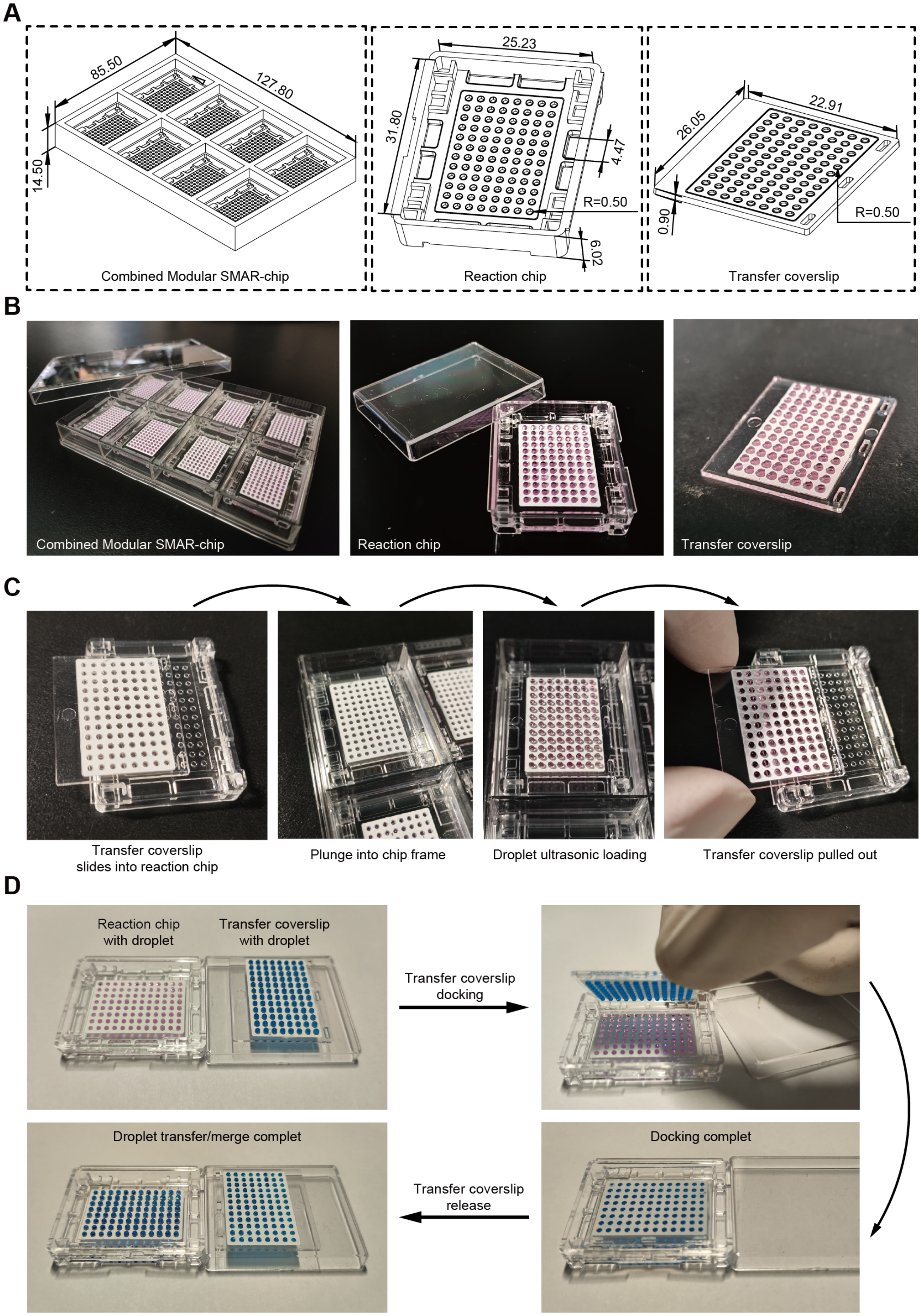


24

**Fig. S1 .Architecture and operation of MoSMAR-chip. (A and B)** Schematic diogram **(A)** and phtographs **(B)** showing the structure and dimensions of the assembled MoSMAR-chip, the reaction chip, and the transfer coverslip. **(C and D)** Images showing the the “spot-cover” procedure of the transfer coverslip pre-loading **(C)** and docking with the reaction chip **(D)** for reagent delivery.

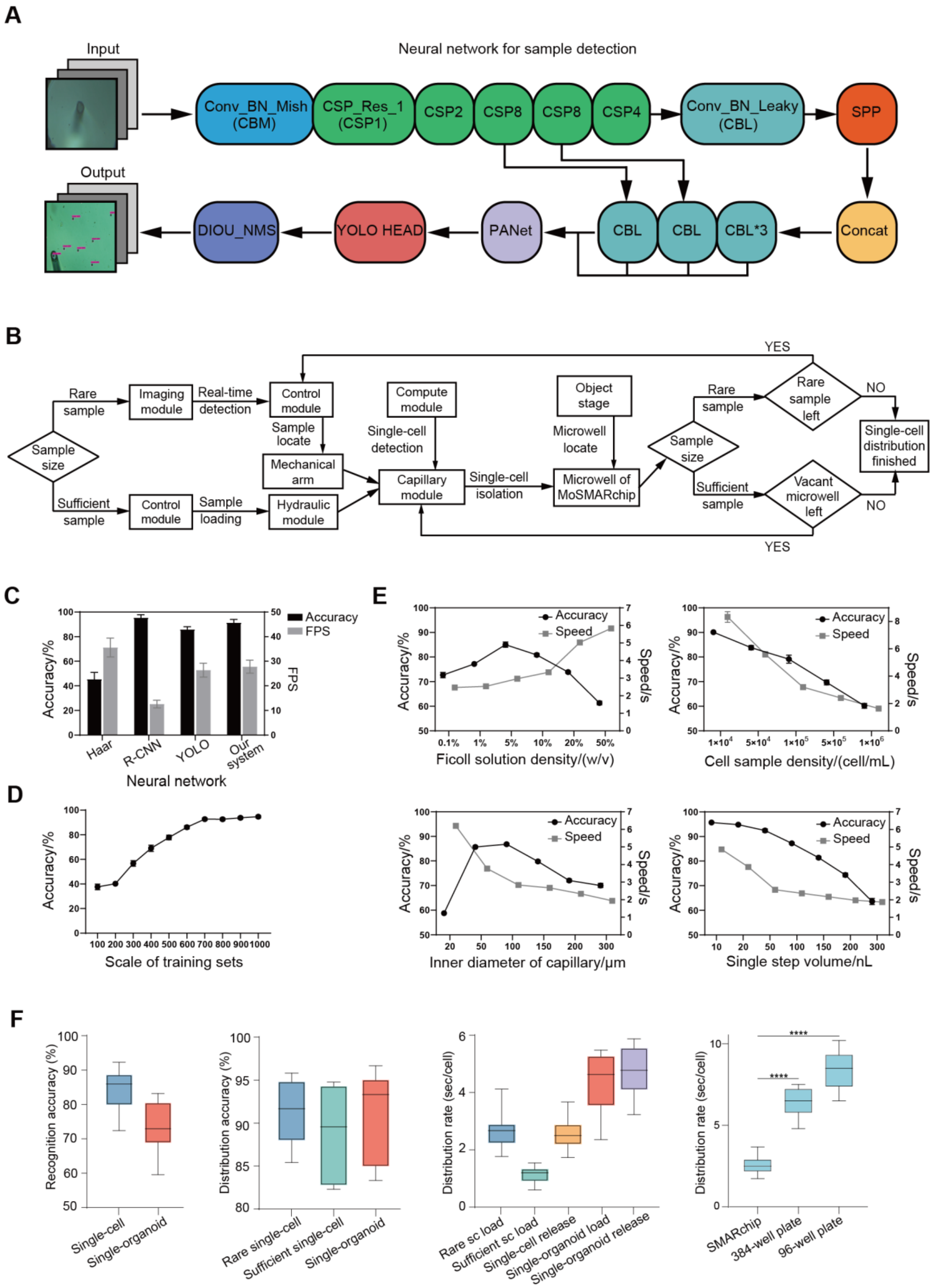
A
Input
Neural network for sample detection
Conv_BN_Mish (CBM)
CSP_Res_1 (CSP1)
CSP2
CSP8
CSP8
CSP4
Conv_BN_Leaky (CBL)
SPP
Output
DIOU_NMS
YOLO HEAD
PANet
CBL
CBL
CBL*3
Concat
B
Sample size
Rare sample
Imaging module
Real-time detection
Control module
Sample locate
Mechanical arm
Sufficient sample
Control module
Sample loading
Hydraulic module
Compute module
Single-cell detection
Capillary module
Single-cell isolation
Object stage
Microwell locate
Microwell of MoSMARchip
Sample size
Rare sample
Rare sample left
YES
NO
Single-cell distribution finished
Sufficient sample
Vacant microwell left
NO
YES
C
Accuracy/%
FPS
Accuracy
FPS
Haar
R-CNN
YOLO
Our system
Neural network
D
Accuracy/%
Scale of training sets
E
Accuracy/%
Speed/s
Accuracy
Speed
Ficoll solution density/(w/v)
Cell sample density/(cell/mL)
Inner diameter of capillary/μm
Single step volume/nL
F
Recognition accuracy (%)
Single-cell
Single-organoid
Distribution accuracy (%)
Rare single-cell
Sufficient single-cell
Single-organoid
Distribution rate (sec/cell)
Rare sc load
Sufficient sc load
Single-cell release
Single-organoid load
Single-organoid release
Distribution rate (sec/cell)
SMARchip
384-well plate
96-well plate
****
****

29

**Fig. S2 .Supplementary illustration and optimization of the automatic single cell distribution instrument. (A)** The major composition of YOLOv4-based neural network classifier adapted in our system. **(B)** The operation logic map of automated single-cell isolation system. **(C to E)** Optimization of the single cell distribution instrument addressed with major influence factors, including neural network type **(C)**, sample scale of neural network training set **(D)**, Ficoll density for cell buffer, cell sample density, inner diameter of capillary, and single step accuracy of hydraulic drive control **(E)**. **(F)** Quantification of the single-cell/organoid recognition accuracy, distribution accuracy, distribution rate under various circumstance, and comparison of the single-cell distribution rates on the MoSMAR-chip and multi-well plates. The center line represents the median value. The bounds of box represent the median values of the upper half and the lower half. The bounds of whiskers represent the maxima and the minima. ****$P<0.0001$, two-sided, unpaired student's *t* tests.

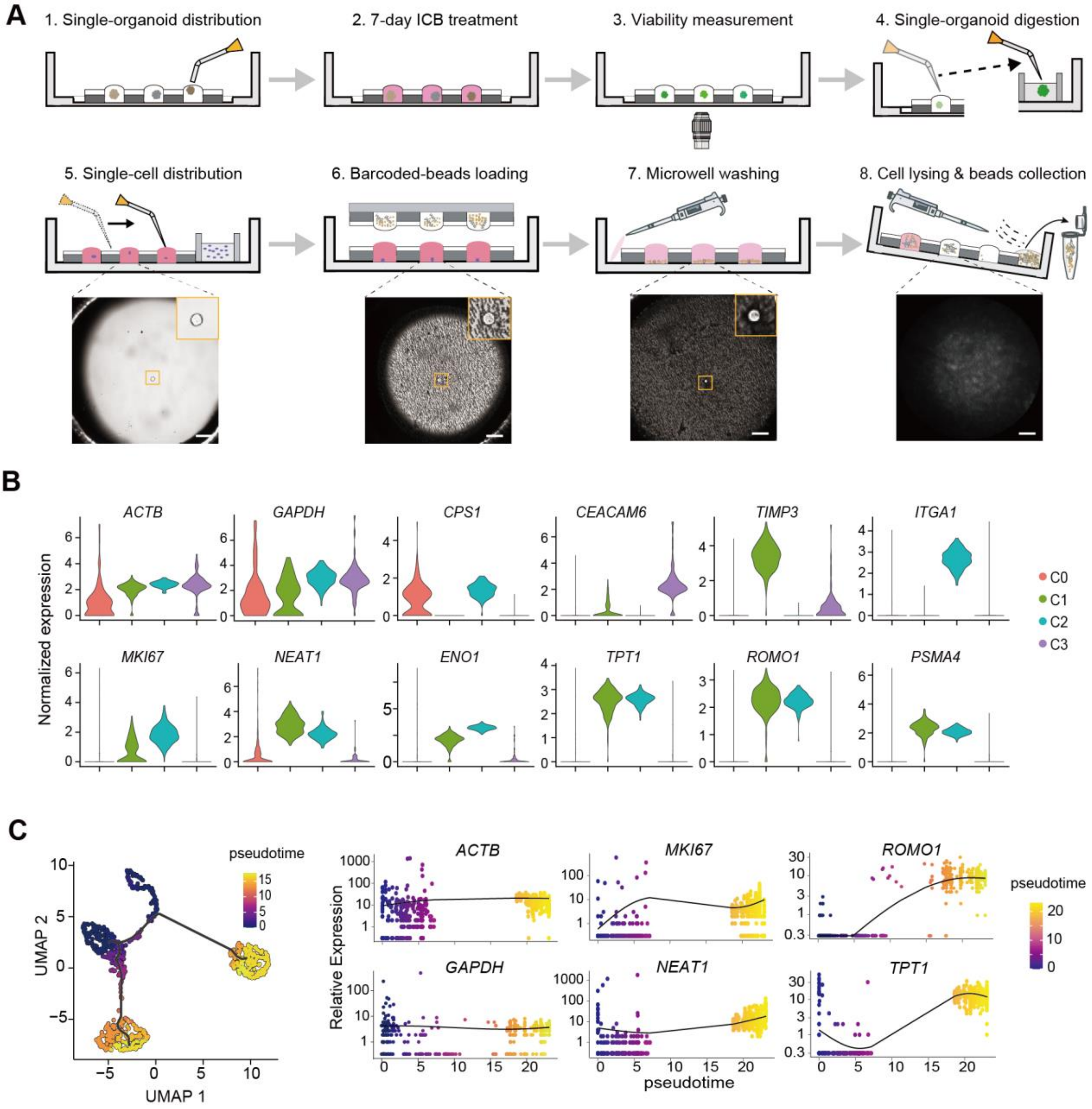


**Fig. S3 .Extended data of Fig. 2**. **(A)** Scheme of the FascRNA-seq procedure for single organoid and images showing single cells in the microwells at the indicated steps. Scale bars, 50 μm. **(B)** Violin plots of gene expression features of the 4 unsupervised UMAP clusters in Fig. 2j. **(C)** Pseudo-trajectory and gene expression analysis along the inferred trajectory.

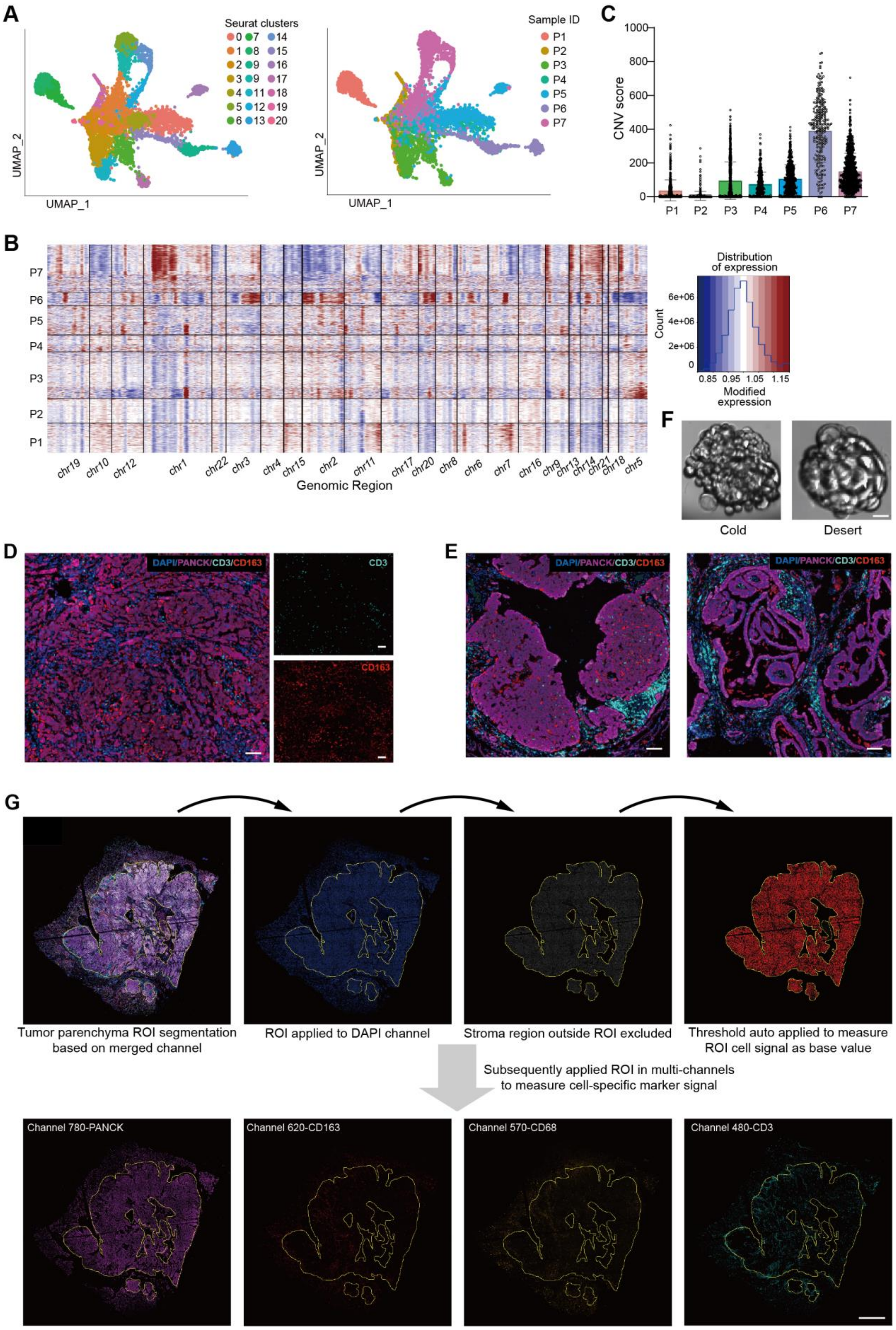

A
Seurat clusters
UMAP_1
UMAP_2
Sample ID
P1
P2
P3
P4
P5
P6
P7
C
CNV score
B
Genomic Region
Distribution of expression
Count
Modified expression
F
Cold
Desert
D
DAPI/PANCK/CD3/CD163
CD3
CD163
E
G
Tumor parenchyma ROI segmentation based on merged channel
ROI applied to DAPI channel
Stroma region outside ROI excluded
Threshold auto applied to measure ROI cell signal as base value
Subsequently applied ROI in multi-channels to measure cell-specific marker signal
Channel 780-PANCK
Channel 620-CD163
Channel 570-CD68
Channel 480-CD3


46

**Fig. S4 .Extended data of Fig 3. (A)** UMAP projections of the 6240 single cells derived from 171 single pLCOs color labeled by unsupervised Seurat clusters (left) or Sample IDs (right). **(B)** Heatmap of the CNV patterns of all the single epithelial cells. Red means amplification and blue indicates deletion. The line chart on the right shows the distribution characteristics of CNV circumstances across single epithelial cells. **(C)** Quantitative assessment of the CNV scores of all the single epithelial cells in pLCOs derived from the 7 patient samples. **(D and E)** IHC images of tumor tissue sections of P1 **(D)** and P4 **(E)** with antibodies specific to T cells (CD3), macrophages (CD163), and epithelial cells (PanCK). Cell nucleus was counterstained with DAPI. scale bars, 100 µm. **(F)** Representative images of the "cold" (left) and "desert" (right) pLCOs derived from P4. Scale bar, 20 µm. **(G)** Workflow of IHC image quantitative analysis for the signals derived from specific cell types in the tumor parenchyma. Scale bar, 1 cm.

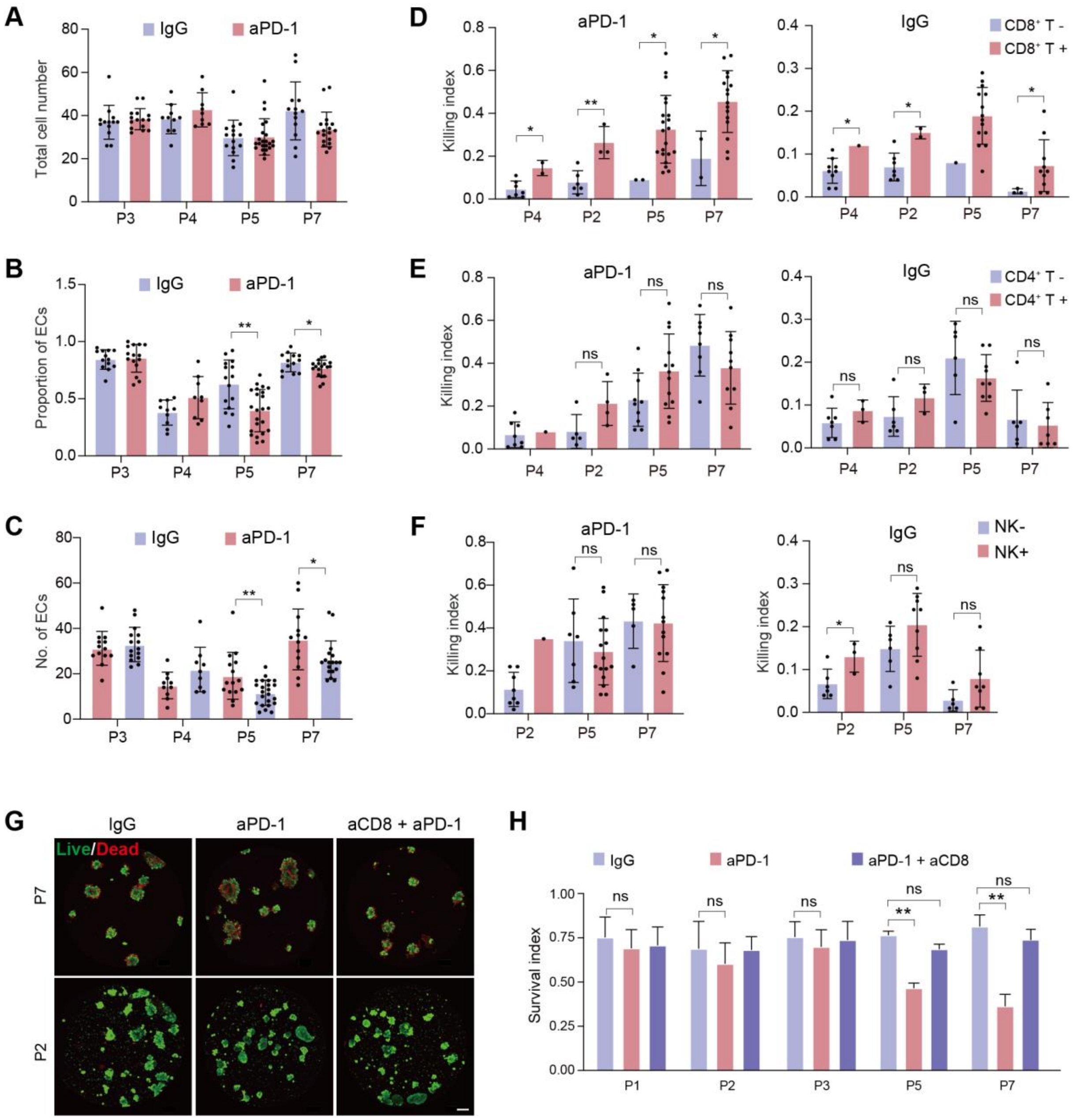


**Fig. S5. Supplementary data demonstrating CD8⁺ T cells mediate the aPD-1 induced tumor cell death.** **(A)** Comparison of total cell numbers in individual pLCOs with or without aPD-1 treatment. Data from 2 aPD-1 sensitive samples (P5 and P7) and 2 insensitive samples (P3, P4) are shown. **(B and C)** Comparisons of the numbers **(B)** and the proportions **(C)** of epithelial cells in single pLCOs with or without aPD-1 treatment demonstrate the significant reduction of epithelial cells in P5 and P7 pLCOs. **(D to F)** Bar graph showing the impact of CD8⁺ T cells **(D)**, CD4⁺ T cells **(E)**, and NK cells **(F)** on aPD-1 induced cell death. **(G)** Images of P2 and P7 pLCOs cultured in 96-well plates under different treatment conditions. Organoids were stained with Calcein AM/PI to indicate the living and dead cells. Scale bar, 100 µm. **(H)** Comparison of the overall organoid viability under different treatment conditions. Note anti-CD8 (aCD8) antibody treatment rescue the cells from aPD-1 induced cell death. **** P<0.0001, paired student's *t* test

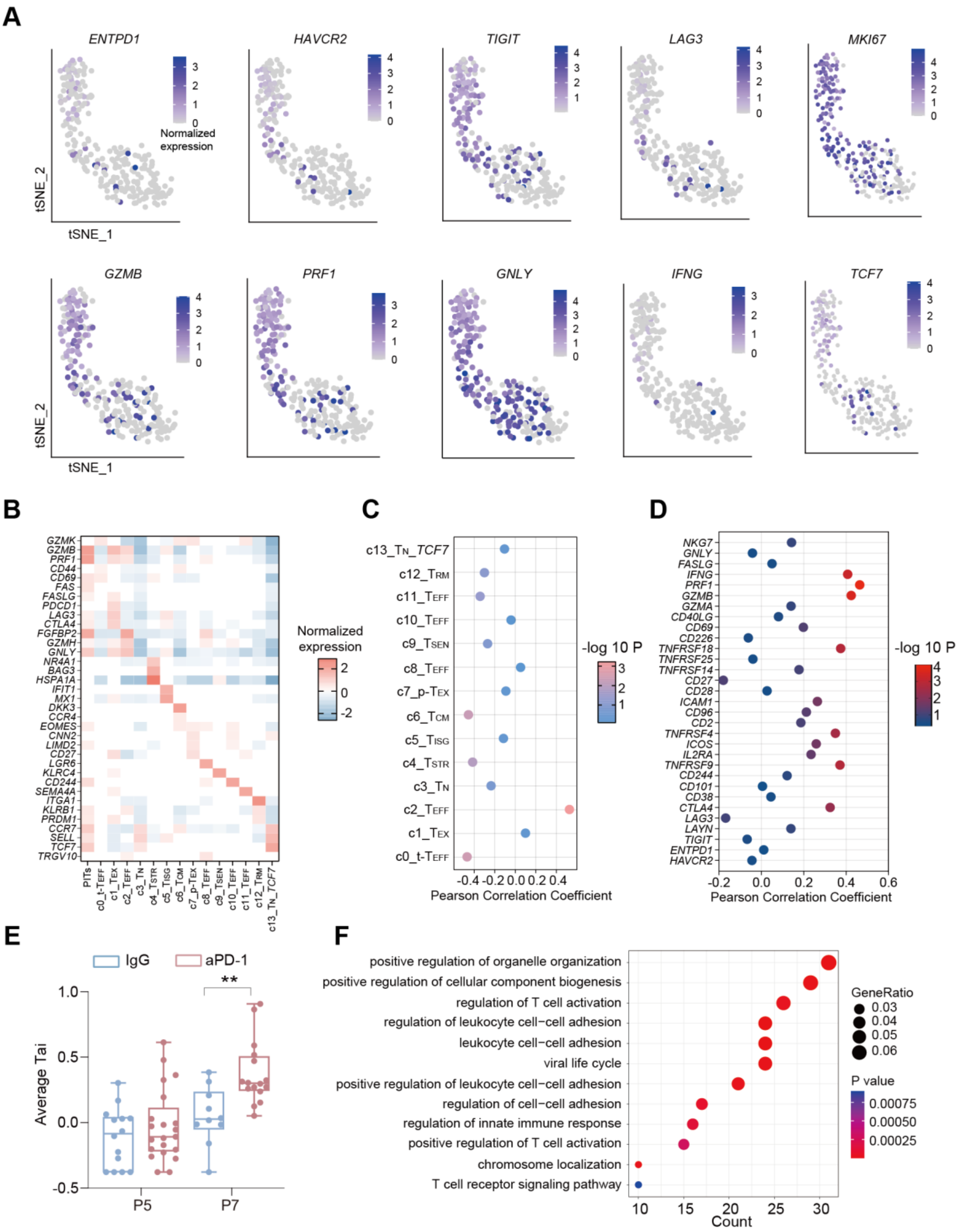

A
ENTPD1
HAVCR2
TIGIT
LAG3
MKI67
Normalized expression
tSNE_2
tSNE_1
GZMB
PRF1
GNLY
IFNG
TCF7
B
Normalized expression
C
Pearson Correlation Coefficient
-log 10 P
D
Pearson Correlation Coefficient
-log 10 P
E
IgG
aPD-1
Average Tai
P5
P7
F
positive regulation of organelle organization
positive regulation of cellular component biogenesis
regulation of T cell activation
regulation of leukocyte cell−cell adhesion
leukocyte cell−cell adhesion
viral life cycle
positive regulation of leukocyte cell−cell adhesion
regulation of cell−cell adhesion
regulation of innate immune response
positive regulation of T cell activation
chromosome localization
T cell receptor signaling pathway
Count
GeneRatio
P value


68

**Fig. S6. Characterization of parenchyma infiltrating CD8$^+$ Ts (PITs) derived from pLCOs. (A)** t-SNE projection of 191 single T cells derived from pLCOs with featureplot of T cell function-associated genes. **(B)** Heatmap illustrating the average expression of T cell function-associated genes in the 191 CD8$^+$ T cells and the 14 CD8$^+$ T clusters reported in a pan-cancer T cell atlas(*23*). **(C)** PCC between the PITs and the CD8$^+$ T clusters in the pan-cancer atlas. **(D)** PCC between the average level of function related genes for CD8$^+$ Ts in a single pLCO and the killing index. 10 genes with good correlation were chosen as a gene set to calculate T cell activation index (Tai). **(E)** Comparison of average Tai in single pLCO under the two treatment conditions. The center line in the box plot represents the median value. The bounds of box represent the median values of the upper half and the lower half. The bounds of whiskers represent the maxima and the minima. $^{**}P < 0.01$, unpaired, two-sided student's *t* test. **(F)** Gene ontology (GO) enrichment analysis of the DEGs between the Tai high (>0.5) and other CD8$^+$ T cells.

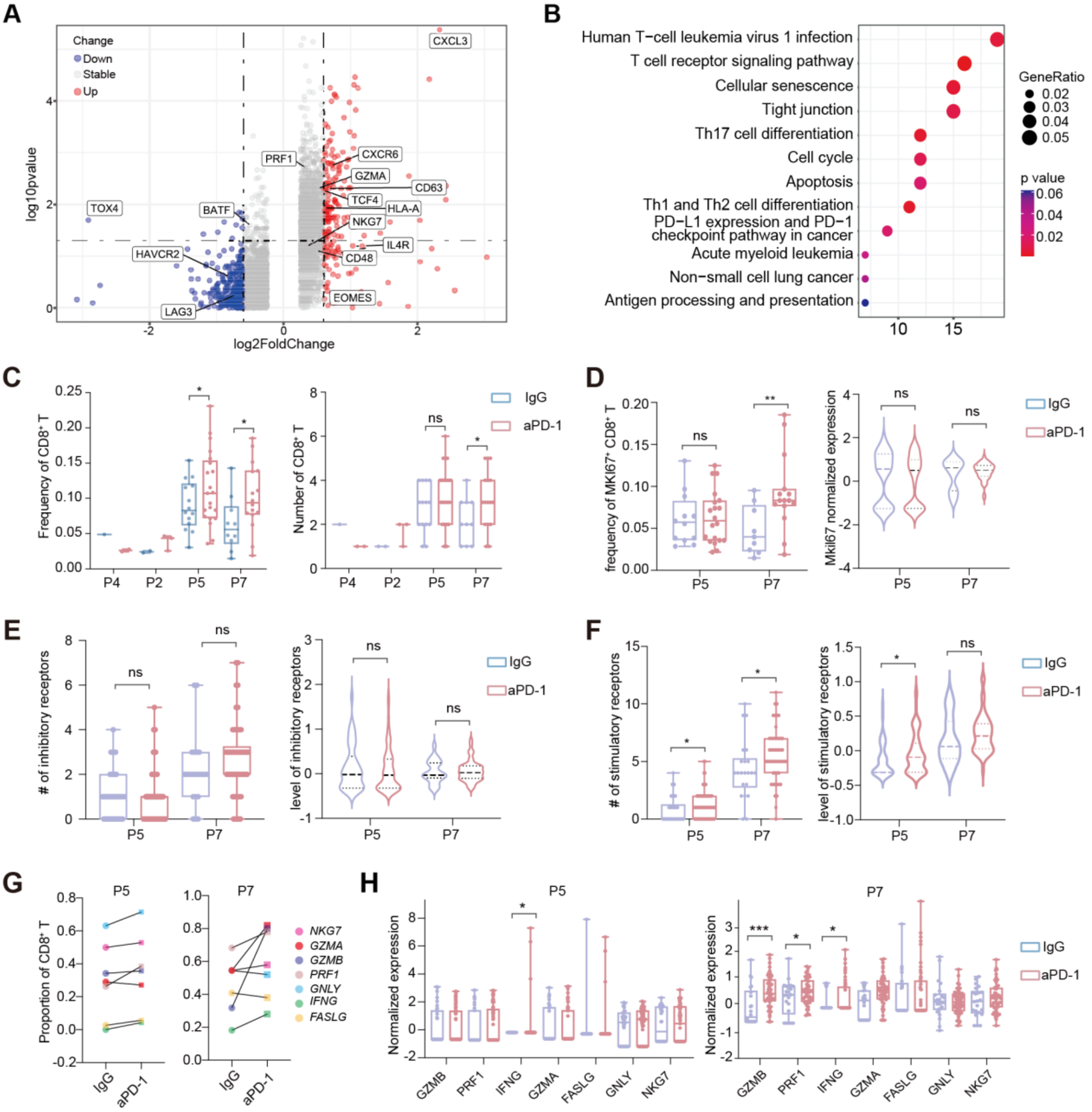

A
Change
Down
Stable
Up
log10pvalue
log2FoldChange
CXCL3
TOX4
BATF
PRF1
CXCR6
GZMA
CD63
TCF4
HLA-A
NKG7
IL4R
CD48
HAVCR2
LAG3
EOMES
B
Human T−cell leukemia virus 1 infection
T cell receptor signaling pathway
Cellular senescence
Tight junction
Th17 cell differentiation
Cell cycle
Apoptosis
Th1 and Th2 cell differentiation
PD−L1 expression and PD−1 checkpoint pathway in cancer
Acute myeloid leukemia
Non−small cell lung cancer
Antigen processing and presentation
GeneRatio
p value
C
Frequency of CD8⁺ T
Number of CD8⁺ T
IgG
aPD-1
P4
P2
P5
P7
D
frequency of MKI67⁺ CD8⁺ T
Mki67 normalized expression
E
### of inhibitory receptors
level of inhibitory receptors
F
### of stimulatory receptors
level of stimulatory receptors
G
Proportion of CD8⁺ T
NKG7
GZMA
GZMB
PRF1
GNLY
IFNG
FASLG
H
Normalized expression
GZMB
PRF1
IFNG
GZMA
FASLG
GNLY
NKG7

**Fig. S7. aPD-1 induced gene expression changes in the $CD8^+$ Ts derived from pLCOs. (A)** Volcano plot showing differentially expressed genes (DEGs) between IgG and aPD-1 treated $CD8^+$ Ts (marked genes with P value < 0.05 (two-sided student's *t* test) and fold change ≥ 1.2). **(B)** Gene ontology (GO) enrichment analysis of the DEGs in (**A**). **(C)** Box plots showing the significant increases in $CD8^+$ T cells in individual pLCOs with the aPD-1 treatment compared to the control IgG. **(D)** Comparison of *KI67* expression in $CD8^+$ Ts under the two treatment conditions. **(E to F)** Expression of the co-stimulatory receptors (*TNFRSF9, IL2RA, ICOS, TNFRSF4, CD2, CD96, ICAM1, CD28, CD27, TNFRSF14, TNFRSF25, TNFRSF18, CD226, CD69,* and *CD40LG*) and the inhibitory receptors (*HAVCR2, ENTPD1, TIGIT, LAYN, LAG3, CTLA4, CD38, CD101 and CD244*) in $CD8^+$ Ts under the two treatment groups. **(G)** Proportions of $CD8^+$ Ts expressing the indicated effector molecules in P5 and P7 pLCOs. **(H)** Comparison of the expression levels of effector molecules in $CD8^+$ T cells under the two treatment conditions. The center lines in the box plots of **(C, D, E, F, and H)** represent the median values. The bounds of box represent the median values of the upper half and the lower half. The bounds of whiskers represent the maxima and the minima. P values were determined by unpaired, two-sided student's *t* test, *P < 0.05, **P < 0.01, ***P < 0.001.

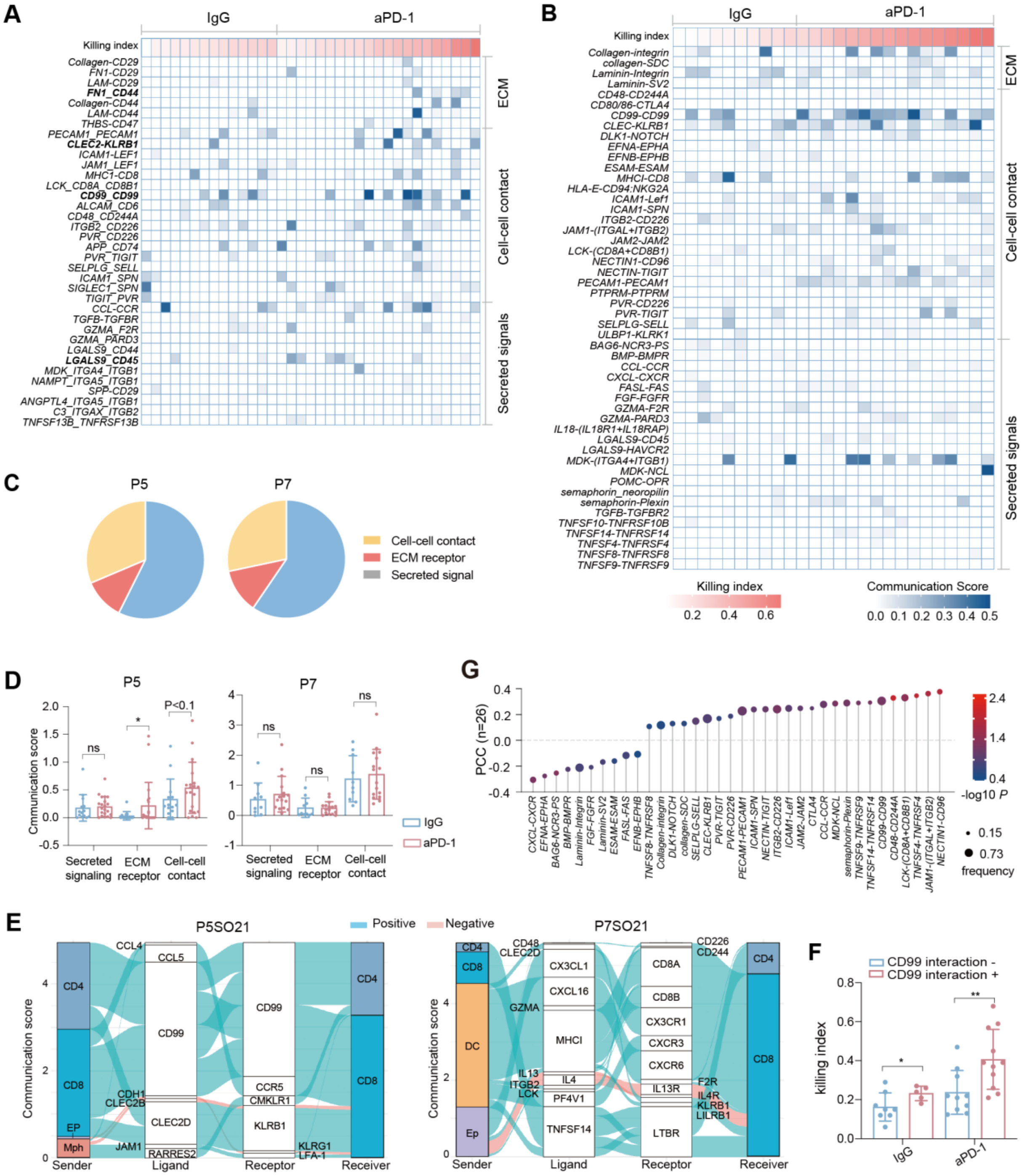
A
IgG
aPD-1
Killing index
Collagen-CD29
FN1-CD29
LAM-CD29
FN1_CD44
Collagen-CD44
LAM-CD44
THBS-CD47
PECAM1_PECAM1
CLEC2-KLRB1
ICAM1-LEF1
JAM1_LEF1
MHC1-CD8
LCK_CD8A_CD8B1
CD99_CD99
ALCAM_CD6
CD48_CD244A
ITGB2_CD226
PVR_CD226
APP_CD74
PVR_TIGIT
SELPLG_SELL
ICAM1_SPN
SIGLEC1_SPN
TIGIT_PVR
CCL-CCR
TGFB-TGFBR
GZMA_F2R
GZMA_PARD3
LGALS9_CD44
LGALS9_CD45
MDK_ITGA4_ITGB1
NAMPT_ITGA5_ITGB1
SPP-CD29
ANGPTL4_ITGA5_ITGB1
C3_ITGAX_ITGB2
TNFSF13B_TNFRSF13B
ECM
Cell-cell contact
Secreted signals
B
IgG
aPD-1
Killing index
Collagen-integrin
collagen-SDC
Laminin-Integrin
Laminin-SV2
CD48-CD244A
CD80/86-CTLA4
CD99-CD99
CLEC-KLRB1
DLK1-NOTCH
EFNA-EPHA
EFNB-EPHB
ESAM-ESAM
MHCI-CD8
HLA-E-CD94:NKG2A
ICAM1-Lef1
ICAM1-SPN
ITGB2-CD226
JAM1-(ITGAL+ITGB2)
JAM2-JAM2
LCK-(CD8A+CD8B1)
NECTIN1-CD96
NECTIN-TIGIT
PECAM1-PECAM1
PTPRM-PTPRM
PVR-CD226
PVR-TIGIT
SELPLG-SELL
ULBP1-KLRK1
BAG6-NCR3-PS
BMP-BMPR
CCL-CCR
CXCL-CXCR
FASL-FAS
FGF-FGFR
GZMA-F2R
GZMA-PARD3
IL18-(IL18R1+IL18RAP)
LGALS9-CD45
LGALS9-HAVCR2
MDK-(ITGA4+ITGB1)
MDK-NCL
POMC-OPR
semaphorin_neoropilin
semaphorin-Plexin
TGFB-TGFBR2
TNFSF10-TNFRSF10B
TNFSF14-TNFRSF14
TNFSF4-TNFRSF4
TNFSF8-TNFRSF8
TNFSF9-TNFRSF9
ECM
Cell-cell contact
Secreted signals
Killing index
0.2 0.4 0.6
Communication Score
0.0 0.1 0.2 0.3 0.4 0.5
C
P5
P7
Cell-cell contact
ECM receptor
Secreted signal
D
P5
P7
Cmmunication score
ns
*
P<0.1
ns
ns
ns
Secreted signaling
ECM receptor
Cell-cell contact
IgG
aPD-1
G
PCC (n=26)
-log10 P
2.4
1.4
0.4
0.15
0.73
frequency
CXCL-CXCR
EFNA-EPHA
BAG6-NCR3-PS
BMP-BMPR
Laminin-Integrin
FGF-FGFR
Laminin-SV2
ESAM-ESAM
FASL-FAS
EFNB-EPHB
TNFSF8-TNFRSF8
Collagen-integrin
DLK1-NOTCH
collagen-SDC
SELPLG-SELL
CLEC-KLRB1
PVR-TIGIT
PVR-CD226
PECAM1-PECAM1
ICAM1-SPN
NECTIN-TIGIT
ITGB2-CD226
ICAM1-Lef1
JAM2-JAM2
CTLA4
CCL-CCR
MDK-NCL
semaphorin-Plexin
TNFSF9-TNFRSF9
TNFSF14-TNFRSF14
CD99-CD99
CD48-CD244A
LCK-(CD8A+CD8B1)
TNFSF4-TNFRSF4
JAM1-(ITGAL+ITGB2)
NECTIN1-CD96
E
P5SO21
Positive
Negative
P7SO21
Communication score
Sender
Ligand
Receptor
Receiver
CD4
CD8
EP
Mph
CCL4
CCL5
CD99
CDH1
CLEC2B
CLEC2D
JAM1
RARRES2
CCR5
CMKLR1
KLRB1
KLRG1
LFA-1
CD48
CLEC2D
GZMA
IL13
ITGB2
LCK
DC
Ep
CX3CL1
CXCL16
MHCI
IL4
PF4V1
TNFSF14
CD226
CD244
CD8A
CD8B
CX3CR1
CXCR3
CXCR6
IL13R
F2R
IL4R
KLRB1
LILRB1
LTBR
F
CD99 interaction -
CD99 interaction +
killing index
*
**
IgG
aPD-1

**Fig. S8. Cell-cell interactions received by $CD8^+$ Ts in individual organoids. (A and B)** Heatmaps showing the cell-cell interactions in individual pLCOs received by $CD8^+$ T cells in P5 **(A)** and P7 **(B)**. The heatmaps are color labeled by the possibility of interactions (i.e., communication score) calculated by CellChat. The bars on the top represent the killing index of individual organoids. **(C)** Pie chart showing the abundance of the three types of interactions. **(D)** Comparison of the communication scores in individual organoids under the two treatment conditions. **(E)** Sankey plots of intercellular communications between various cell types and T cells in P5SO21 and P7SO21 organoids. The positive or negative color label of ligand-receptor connect were determined by background knowledge. **(F)** T cells receiving CD99 homophilic interaction responded to aPD-1 more significantly. **(G)** Analysis of Pearson correlation between communication scores of ligand-receptor pairs and corresponding killing index of P7 pLCOs. P values were determined by unpaired, two-sided student's *t* test, $*P < 0.05$, $**P < 0.01$, $***P < 0.001$.

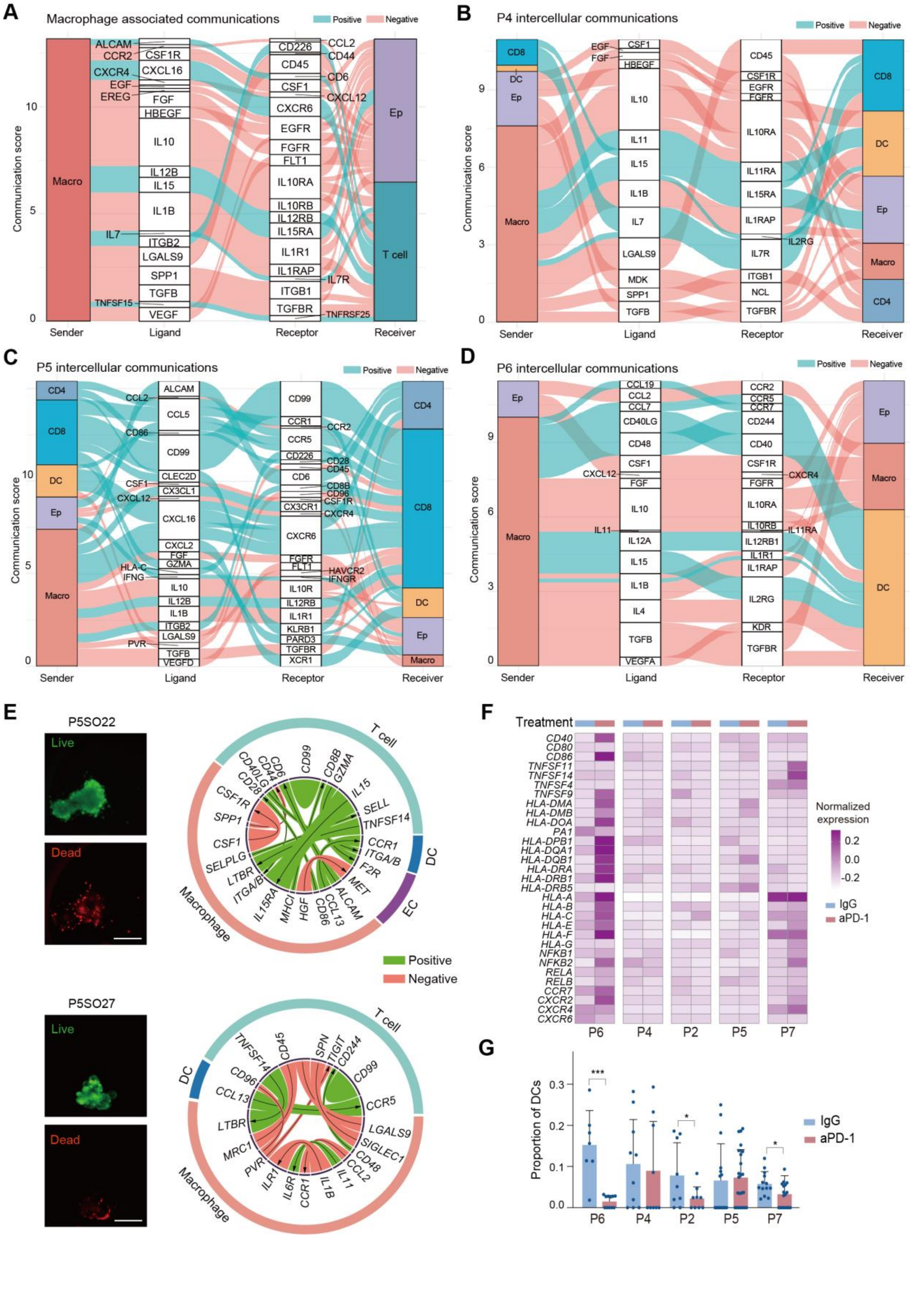
A
Macrophage associated communications
Positive
Negative
Communication score
Sender
Ligand
Receptor
Receiver
Macro
Ep
T cell
ALCAM
CCR2
CXCR4
EGF
EREG
IL7
TNFSF15
CSF1R
CXCL16
FGF
HBEGF
IL10
IL12B
IL15
IL1B
ITGB2
LGALS9
SPP1
TGFB
VEGF
CD226
CD45
CSF1
CXCR6
EGFR
FGFR
FLT1
IL10RA
IL10RB
IL12RB
IL15RA
IL1R1
IL1RAP
ITGB1
TGFBR
CCL2
CD44
CD6
CXCL12
IL7R
TNFRSF25
B
P4 intercellular communications
CD8
DC
Ep
Macro
CD4
EGF
FGF
CSF1
HBEGF
IL10
IL11
IL15
IL1B
IL7
LGALS9
MDK
SPP1
TGFB
CD45
CSF1R
EGFR
FGFR
IL10RA
IL11RA
IL15RA
IL1RAP
IL2RG
IL7R
ITGB1
NCL
TGFBR
C
P5 intercellular communications
CD4
CD8
DC
Ep
Macro
CCL2
CD86
CSF1
CXCL12
HLA-C
IFNG
PVR
ALCAM
CCL5
CD99
CLEC2D
CX3CL1
CXCL16
CXCL2
FGF
GZMA
IL10
IL12B
IL1B
ITGB2
LGALS9
TGFB
VEGFD
CCR1
CCR5
CD226
CD6
CX3CR1
CXCR6
FGFR
FLT1
IL10R
IL12RB
IL1R1
KLRB1
PARD3
TGFBR
XCR1
CCR2
CD28
CD45
CD8B
CD96
CSF1R
CXCR4
HAVCR2
IFNGR
D
P6 intercellular communications
CCL19
CCL2
CCL7
CD40LG
CD48
CSF1
CXCL12
FGF
IL10
IL11
IL12A
IL15
IL1B
IL4
TGFB
VEGFA
CCR2
CCR5
CCR7
CD244
CD40
CSF1R
FGFR
IL10RA
IL10RB
IL12RB1
IL1R1
IL1RAP
IL2RG
KDR
TGFBR
CXCR4
IL11RA
E
P5SO22
Live
Dead
T cell
DC
EC
Macrophage
Positive
Negative
P5SO27
Live
Dead
F
Treatment
CD40
CD80
CD86
TNFSF11
TNFSF14
TNFSF4
TNFSF9
HLA-DMA
HLA-DMB
HLA-DOA
PA1
HLA-DPB1
HLA-DQA1
HLA-DQB1
HLA-DRA
HLA-DRB1
HLA-DRB5
HLA-A
HLA-B
HLA-C
HLA-E
HLA-F
HLA-G
NFKB1
NFKB2
RELA
RELB
CCR7
CXCR2
CXCR4
CXCR6
P6
P4
P2
P5
P7
Normalized expression
0.2
0.0
-0.2
IgG
aPD-1
G
Proportion of DCs
***
*
*
IgG
aPD-1

**Fig. S9. Specific features of TIME for individual patients. (A)** Sankey plot showing cellular interaction between Mphs and ECs or T cells. Cells from all the P4 and P5 organoids were pooled together for the analysis. **(B to D)** Sankey plots of intercellular communication landscapes in P4 **(B)**, P5 **(C)**, and P6 **(D)**. Cells from the same tumor sample were pooled together for the analysis. **(E)** Images and interactive circus plots of organoids P5SO22 and P5SO27. Scale bars, 100 μm. **(F)** Heatmap showing the average of the normalized expression of DC maturation related genes under the two treatment conditions. **(G)** Comparison of DC proportion in individual organoids under the two treatment conditions. P values were determined by unpaired, two-sided student's *t* test, $^{*}P < 0.05$, $^{***}P < 0.001$.

#### Supplementary Table Legends

Table S1. Top 40 GO terms most significantly enriched in C3 of P5

Table S2. Materials of home-made superhydrophobic paint

Table S3. Components of automated single-cell distribution instrument

Table S4. Reagents and kits used in FascRNA-seq

Table S5. The primers and barcode sequences of FascRNA-seq

Table S6. The composition of the LCOM medium

Table S7. Reagents and kits used for fresh tumor/ tumor organoids treatment

#### Description for Movies S1 to S3

Movie S1. The process of single-cell loading and distribution based on SCDI.

Movie S2. The process of single-organoid loading and distribution based on SCDI.

Movie S3. The horizontal comparison of SCDI operation on MoSMARchip and a 384well-plate.
